# UniversalEPI: robust prediction of cell type-specific and differential chromatin interactions from DNA sequence and chromatin accessibility

**DOI:** 10.1101/2024.11.22.624813

**Authors:** Aayush Grover, Lin Zhang, Till Muser, Simeon Häfliger, Minjia Wang, Josephine Yates, Marie-Claire Indilewitsch, Eliezer M. Van Allen, Fabian J. Theis, Ignacio L. Ibarra, Ekaterina Krymova, Valentina Boeva

## Abstract

Enhancer–promoter interactions (EPIs) play a central role in gene regulation, but experimental techniques such as Hi-C for mapping these interactions remain costly and labor-intensive. Computational methods have been developed to predict EPIs in silico from DNA sequence and chromatin information; however, there are major challenges with the generalizability and accuracy of predictions by existing methods across cell types and conditions unseen during model training. We developed and validated UniversalEPI, an attention-based deep ensemble model that predicts EPIs up to 2 Mb apart using only DNA sequence and chromatin accessibility (ATAC-seq) data. Unlike models that reconstruct full Hi-C contact maps, UniversalEPI focuses on biologically relevant, sparse chromatin interactions between accessible regulatory elements. It generalizes across both bulk and single-cell ATAC-seq-derived pseudo-bulk datasets, delivering state-of-the-art performance while using fewer input modalities than existing approaches. By modeling predictive uncertainty, UniversalEPI enables statistically robust differential analysis of chromatin interactions across conditions. We demonstrate its utility by tracking dynamic EPIs during human macrophage activation and identifying regulatory differences between cancer cell states in esophageal adenocarcinoma. By providing precalculated Hi-C predictions for 157 ENCODE datasets, UniversalEPI expands the scope and applicability of in silico 3D genome modeling for studying gene regulation in development and disease.

## 1 Introduction

In complex organisms, different cell types have distinct transcriptional programs that lead to their specific functions. Chromatin interactions between gene promoters and cis-regulatory elements, specifically enhancers, modulate gene expression through the formation of three-dimensional chromatin architecture. These interactions facilitate the recruitment of ubiquitously expressed or tissue-specific transcription factors (TFs), coactivators, and the basal transcription machinery to gene promoters, enabling precise spatial and temporal control of gene expression^1^. Enhancer-promoter looping is mediated by protein complexes such as YY1, CTCF, cohesin, and mediator, which help establish and stabilize these chromatin contacts. CTCF facilitates long-range chromatin interactions by organizing topologically associating domains (TADs), which compartmentalize the genome into functionally distinct regions with an increased probability of interactions between enhancers and promoters within the same TAD^2,3^. TADs are typically bordered by the CTCF protein bound in the convergent orientation (i.e., forward and reverse CTCF motifs are found enriched at interacting TAD boundaries)^4^. Additionally, the zinc-finger TF YY1 acts as a structural tether by forming DNA loops that bring enhancers and promoters into close proximity^5^. Moreover, there are several lines of evidence that the TF SP1 contributes to chromatin architecture^6–8^. SP1 binds to GC-rich sequences at promoters and enhancers, increasing chromatin accessibility.

The dynamic nature of enhancer–promoter interactions (EPIs) allows cells to respond to developmental cues, environmental signals, and other regulatory inputs, ensuring appropriate gene expression that is necessary for cellular identity and function^9,10^. Disruptions in chromatin interactions, such as through mutations or chromosome instability, are associated with various genetic diseases, including cancer^11,12^. Thus, studying how chromatin interactions influence gene expression is crucial for understanding cell differentiation, development, and disease. Chromatin interactions can be profiled experimentally using Hi-C, an unbiased but costly and labor-intensive high-throughput chromosome conformation capture (3C) technique. However, due to the resource demands of Hi-C, interest has increasingly shifted to in silico methods for inferring EPIs and their dynamics in different conditions.

Several recent studies have proposed deep learning-based architectures to predict EPIs. This includes DeepC, Akita, and Orca, which predict chromatin interactions in a specific cell type from the DNA sequence alone^13–15^. These methods can be used to assess the effects of genomic variants on chromatin structure in a cell type of interest. Although they generate realistic Hi-C maps, by design, these methods do not generalize to cell types that are unseen during model training. Methods such as DeepTACT, HiC-Reg, and Epiphany that can generalize and thereby predict cell-type-specific chromatin interactions in cell types that were not used in training require additional epigenetic features as inputs, such as chromatin accessibility and histone post-translational modifications^16–18^. Nevertheless, these methods can potentially overfit to cell types used in the model training because, by construction, they capture interacting DNA motifs corresponding to tissue-specific TFs that are expressed in the training cell types. Moreover, these methods require diverse input data, such as DNA accessibility (ATAC-seq) and chromatin immunoprecipitation followed by sequencing (ChIP-seq) of histone H3 lysine K27 acetylation (H3K27ac), which are not always readily available. Some other methods focus on predicting interactions between a given enhancer and a promoter using only epigenetic features^19,20^. However, such methods cannot be used to understand the mechanisms of non-coding mutations as they cannot capture the effect of mutations in insulators, which modulate EPIs.

The C.Origami and EPCOT methods currently provide state-of-the-art solutions for predicting cell-type-specific chromatin interactions from bulk data^21,22^. Both approaches combine convolutional neural networks (CNNs) with transformer architectures but differ in their required input modalities and predicted outputs. While C.Origami requires DNA sequence, chromatin accessibility (ATAC-seq), and CTCF binding affinity (ChIP-seq), EPCOT utilizes only DNA sequence and chromatin accessibility data. Importantly, EPCOT generates Z-score–normalized contact maps, which do not provide information on absolute interaction frequencies, whereas C.Origami predicts raw Hi-C interaction values, facilitating quantitative comparisons across conditions and more precise interpretation of biologically meaningful differences in chromatin interactions. Due to their complexity and size, C.Origami was initially trained on just one cell line (IMR90), whereas EPCOT was trained using data from four cell lines (H1-hESC, A549, GM12878, and HeLa-S3).

More recently, ChromaFold was introduced to predict chromatin interactions using single-cell ATAC-seq (scATAC-seq) data and occurrences of CTCF motifs derived from the DNA sequence^23^. ChromaFold addresses several limitations of the earlier methods: it eliminates the requirement for CTCF ChIP-seq data, significantly reduces training time, and efficiently incorporates data from multiple cell types. Moreover, ChromaFold is unlikely to overfit to the training cell types because it does not account for any DNA motifs other than those bound by CTCF. The major shortcomings of ChromaFold are that instead of predicting raw Hi-C counts, it estimates distance-and GC-content-based Z-scores, and the method was developed explicitly for single-cell inputs. Thus, there remains a need for a fast and accurate method that can predict raw Hi-C from both single-cell and bulk chromatin accessibility data, can generalize across cell types unseen during training, and does not require additional inputs such as CTCF ChIP-seq. Importantly, to be able to predict statistically significant changes in EPIs across conditions, *e.g.*, variation due to DNA variants or epigenetic differences across transcriptional cell states, the ideal method should output uncertainty values along with predictions.

In this work, we introduce UniversalEPI, an extremely lightweight and accurate deep-ensemble model consisting of CNN layers and transformer blocks. This method overcomes the shortcomings of others by training on automatically extracted DNA binding motifs of three ubiquitously expressed TFs, CTCF, YY1, and SP1^24–26^, complemented with the ATAC-seq profile from DNA accessible regions. This choice of architecture allows for reducing noise and model size while still capturing all functionally relevant interactions in large genomic domains (up to 2Mb). Furthermore, we incorporate uncertainty estimation by integrating both aleatoric (data) and epistemic (model) uncertainty using stochastic training loss^27^ and deep ensemble^28^, enhancing the predictive reliability of the model. Therefore, unlike earlier methods that provide only point estimates, our approach also quantifies prediction confidence. Using the reported maximum-confidence fold-change (FC), a unique feature of UniversalEPI that removes inherent data noise, the user can focus on significantly altered EPIs across conditions.

After training and testing UniversalEPI on four human cell lines with all necessary input data, we benchmarked UniversalEPI against the state-of-the-art methods. We found its performance in predicting chromatin interactions in unseen human cell types was significantly superior to methods such as Akita, EPCOT, C.Origami, and Chromafold. We showed that UniversalEPI can be used to assess chromatin dynamics across conditions by using data generated by Reed *et al.* on human macrophage activation^29^. The uncertainty-aware UniversalEPI predictions agreed with the ground-truth Hi-C measurements with Spearman’s correlation above 0.9. Finally, we showed that UniversalEPI can predict Hi-C interactions from pseudo-bulks of single-cell ATAC-seq data profiles and reveal chromatin dynamics across undifferentiated and differentiated cell states in human esophageal adenocarcinoma, specifically in promoters of genes encoding master transcriptional regulators. Together, these findings show that UniversalEPI is a lightweight, generalizable model that accurately predicts EPIs using bulk or single-cell ATAC-seq and DNA sequencing data in unseen cell types while indicating a level of uncertainty. This model now makes it possible to carry out in silico experiments to assess changes in chromatin folding under diverse biological scenarios.

## 2 Results

### 2.1 UniversalEPI: a transformer-based model to predict genomic interactions from DNA sequence and chromatin accessibility

UniversalEPI is a two-stage deep learning model that comprises two sequentially trained neural networks that use DNA sequences and signals from ATAC-seq from training cell lines as input and ground-truth ChIP-seq and Hi-C data as targets (Figure 1a). To infer the Hi-C signal in a test cell type, UniversalEPI requires DNA sequence and cell-type-specific ATAC-seq profiles as input (Figure 1b).

**Figure 1.**
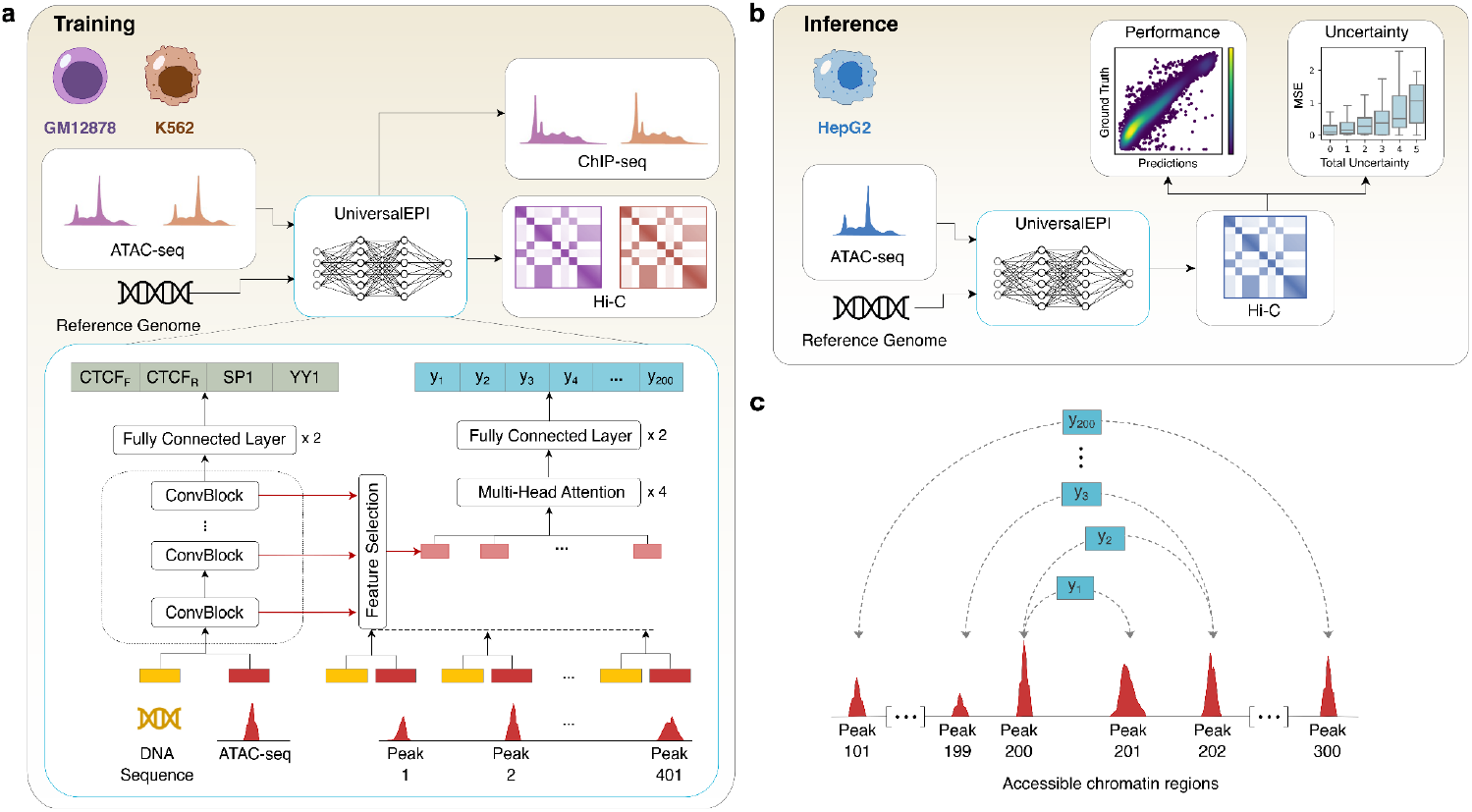
The UniversalEPI model architecture. **a,** UniversalEPI uses one-hot encoded DNA sequence and ATAC-seq p-value tracks as input to predict cell-type-specific transcription factor binding and chromatin interactions. The first stage of UniversalEPI consists of a convolutional architecture that predicts transcription factor (CTCF forward and reverse, SP1 and YY1) binding affinity from DNA sequence and ATAC-seq data. The second stage automatically selects the features from the pre-trained convolutional layers from the first stage using a stochastic gating mechanism. The multi-head attention block then uses these features to predict cell-type-specific Hi-C interactions between accessible regions. **b,** The trained UniversalEPI can then be used to predict Hi-C interactions for an unseen cell type. The model also outputs uncertainty associated with each prediction, which increases with the mean squared error (MSE) of the model. **c,** UniversalEPI predicts pairwise Hi-C interactions (*y*_*i*_) between the central 200 accessible regions corresponding to ATAC-seq peaks.

The first neural network of UniversalEPI was trained to predict the genome-wide binding occupancy of the TFs SP1, CTCF, and YY1 using maximal intensities from ChIP-seq data in human cancerous and non-cancerous cell lines GM12878 and K562 (Model 1) and IMR90 and HepG2 (Model 2). This was achieved by training a one-dimensional 5-layer CNN, producing a 4-dimensional output on the concatenation of one-hot encoded 1Kb DNA sequences and its corresponding ATAC-seq p-value signals (Methods). Since the orientation of CTCF binding to chromatin plays an important role in determining the stability of TADs, we split the CTCF ChIP-seq based on orientation (forward or reverse motifs) and predicted the corresponding maximal ChIP-seq intensities as separate targets. The pre-trained convolutional layers from the first stage thus generated sequence embeddings, indicating the presence or absence of the three TFs in each accessible region of the genome. This design of sequence embeddings was chosen to prevent capturing DNA motifs of other, potentially cell-type-specific, TFs and thereby enable the model to generalize predictions across different cell types at the second stage.

In the second stage, we trained the model to predict Hi-C values between accessible regions based on the DNA sequence embeddings calculated in the first stage (Figure 1c). By focusing on accessible regions, we retain most active regulatory elements, ensuring that the chromatin interactions between key regions involved in gene regulation are predicted by the model (Figure S1). We employed a transformer-based encoder because of its ability to capture long-range interactions and create large-context embeddings of genomic elements (Methods). This allowed the model to efficiently learn the inter-dependency between the input regions and, consequently, predict the Hi-C interactions between regulatory elements with high accuracy in a cell-type-specific manner. Specifically, the model input in the second stage corresponded to 401 consecutive accessible regions (or peaks) spanning in total about 4Mb. Each region was 1Kb long, centered around the position of the maximum ATAC-seq signal. Skipping the information about the DNA sequence between accessible regions allowed for a lightweight architecture of UniversalEPI (2.5M parameters) that can run on standard GPUs while maintaining a large receptive field and making accurate predictions for interactions between regulatory elements that are up to 2Mb apart.

To provide uncertainty estimations and make the model robust to the initialization parameters, we implemented a deep ensemble of 10 single models^28^. This approach allowed the estimation of uncertainty for each prediction, facilitating confident differential predictions between two conditions, *i.e.*, when inputs (DNA or ATAC-seq signal) change. The total predictive uncertainty reported was a combination of two types: aleatoric uncertainty, which stems from inherent data variability, and epistemic uncertainty, which reflects model limitations. The former was estimated by minimizing the negative log-likelihood loss to incorporate the predicted variance, whereas the latter was quantified through the variability across the deep ensemble (Methods).

To ensure the stability of the proposed architecture, we trained both stages of UniversalEPI in a cross-validation setting on human cancerous and non-cancerous cell lines. Model 1 was trained on GM12878 and K562 cells and tested on unseen chromosomes of IMR90 and HepG2 cells. Model 2 was trained on IMR90 and HepG2 cells and tested on unseen chromosomes of GM12878 and K562 cells. Approximately 15% of the human genome was set aside for validation (chromosomes 5, 12, 13, and 21) and approximately 15% for testing (chromosomes 2, 6, and 19). The first stage of UniversalEPI predicted TF binding on unseen chromosomes and unseen cell types for CTCF, YY1, and SP1 with an average Pearson’s correlation of 0.78, 0.62, and 0.47, respectively (Figure S2). We further validated that the first stage of our model learned biologically relevant information in the input DNA sequence by analyzing the model gradients using the DeepLIFT attributions scores^30^. These attribution scores revealed important DNA motifs associated with the binding of each TF, closely resembling their respective known consensus motifs (Figure S2).

### 2.2 UniversalEPI can predict changes in TAD organization in response to in silico modifications

Experimental Hi-C values, which quantify chromatin interactions between each pair of genomic regions, are inversely correlated with the distance between regions and follow the exponential decline resulting from polymer physics^31^. Similarly, UniversalEPI’s single model predictions exponentially declined with distance on the test chromosomes of the unseen cell lines, highlighting the model’s ability to accurately capture the effect of distance on chromatin structure (Figure 2a). Further, to show that UniversalEPI was sensitive to variability in chromatin accessibility and thus is capable of making predictions for unseen cell types based on ATAC-seq inputs, we computed the distance-stratified correlation between the predicted and ground-truth Hi-C signal between pairs of accessible regions (Methods). Using as inputs cell-line-specific ATAC-seq profiles, UniversalEPI obtained equally high values of distance-stratified correlation between predicted and ground-truth Hi-C values for all four test cell lines, IMR90 and HepG2 (Model 1), and GM12878 and K562 (Model 2) (Figure 2b).

**Figure 2.**
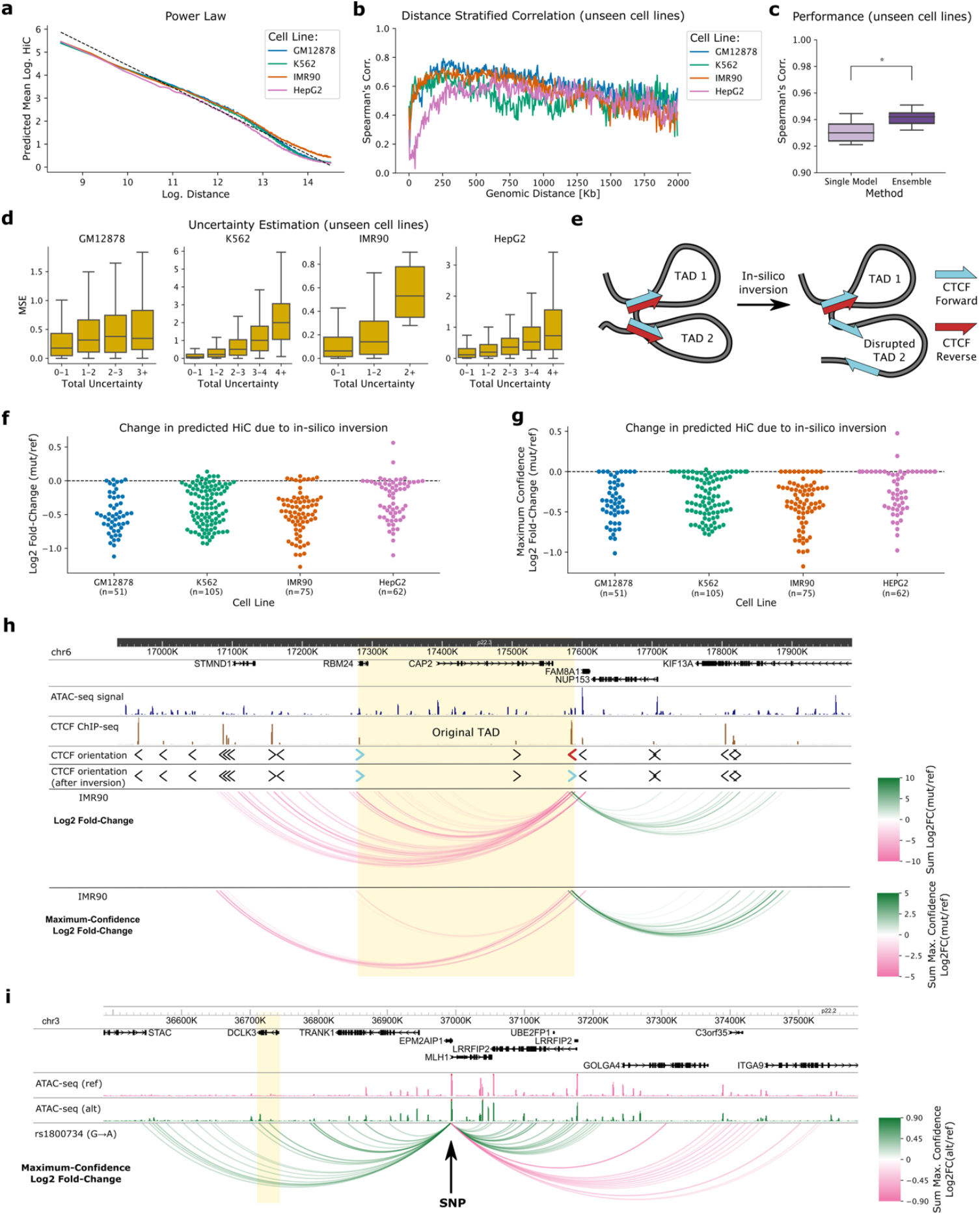
UniversalEPI learns information from biologically relevant variables and provides well-calibrated uncertainty estimates. **a,** The predicted Hi-C signal by UniversalEPI follows the power law resulting from polymer physics. The average predicted signal is shown for test chromosomes of test cell lines (*i.e.*, cell lines unseen during training): GM12878, K562, IMR90, and HepG2. The theoretical relationship between the log-distance and log-Hi-C values is depicted as the dotted line. **b,** Distance-stratified Spearman’s correlation for all the unseen cell lines during training displaying UniversalEPI’s sensitivity to the input ATAC-seq. **c,** Spearman’s correlation is compared between the predictions made by a single model as compared to the ensemble. The predictions are made on the test chromosomes of test cell lines. A Mann-Whitney U test is used to compare the methods; *: P ≤ 0.05. **d,** Relationship between ensemble’s predictive error, calculated as a mean squared error (MSE), and prediction uncertainty is illustrated on the test chromosomes of test cell lines. **e,** Graphical illustration depicting the effect of in-silico inversion of a CTCF binding site on genome organization. **f,** Log2 fold-change between the UniversalEPI predictions before and after the in-silico inversion. The experiment is done for all TADs in test chromosomes in unseen cell lines: GM12878, K562, IMR90, and HepG2. **g,** Maximum-confidence Log2 fold-change between the UniversalEPI predictions before and after the in-silico inversion (Methods), same regions as in (f). **h,** An example of UniversalEPI predictions for a TAD in chromosome 6 of IMR90 cells (unseen cell line and unseen chromosome). The original TAD is highlighted in yellow. The change in Hi-C predicted by an ensemble model is calculated using log2 fold-change before and after the in-silico inversion (top track) and maximum-confidence log2 fold-change (bottom track). In the presence of multiple ATAC-seq peaks within the same 5Kb bin, the log2 fold-change is summed among all pairs of interactions contained in the two bins. **i,** Maximum-confidence log2 fold-change of Hi-C measurements predicted by UniversalEPI for the regulatory element encompassing the single-nucleotide polymorphism (SNP) rs1800734 (G → A) as one of the endpoints. The *DCLK3* gene, experimentally shown to exhibit stronger interactions with the alternate allele (A)^35^, is highlighted in yellow.

Next, we evaluated how the deep ensemble implemented in UniversalEPI improved prediction accuracy compared to a single model. As expected, the deep ensemble approach significantly outperformed the single model in generalizing to test chromosomes from previously unseen cell lines (Figure 2c). Notably, higher total uncertainty was generally associated with a greater mean-squared error between predicted and ground-truth Hi-C values, indicating that the uncertainty estimates were well calibrated (Figure 2d).

We then assessed the sensitivity of the deep ensemble UniversalEPI model to changes in the DNA sequence by implementing in silico inversions of CTCF binding sites located at TAD boundaries. Disruptions in TADs have been linked to diseases such as cancer, making chromatin interactions within and across TADs important for exploring disease mechanisms^32–34^. For 293 TADs across the four cell lines, we inverted one endpoint of a TAD boundary containing a CTCF site in a convergent orientation, which was expected to significantly reduce the strength of chromatin interactions at the original TAD boundary (Figure 2e). Indeed, when we provided the original and modified inputs to UniversalEPI, we predominantly observed negative values of the log FC across all four test cell lines, indicating a decrease in predicted interactions following CTCF motif inversion (Figure 2f). The maximum-confidence FC values, estimated by our uncertainty-aware model to represent the largest changes with 90% probability, aligned with the original predictions (Figure 2g, Methods). This consistency can be attributed to the strong effect of the in silico change to the TAD boundary.

The reported prediction uncertainty played an important role in determining other significantly altered interactions predicted by UniversalEPI after the in-silico inversion of CTCF binding motifs at TAD boundaries. We illustrated this effect using a randomly selected TAD in IMR90 cells (chr6:17,280,000-17,585,000). UniversalEPI predicted a substantial reduction in interaction between the original TAD boundaries as well as between the 3’ TAD boundary and several enhancer regions upstream of the original 5’ TAD boundary (Figure 2h). Moreover, we observed a significant increase in predicted interactions with a reverse CTCF motif located nearly 200Kb away, downstream of the original 3’ TAD boundary. Importantly, the estimated uncertainty allowed the calculation of the maximum-confidence FC and retention of only the highly reliable differential interactions (Figure 2h, Methods). This result confirmed that UniversalEPI is capable of capturing the effects of CTCF binding motif orientation on the stability and organization of TADs, and this is maintained when filtering for significant differential interactions using the estimated prediction uncertainty.

To further evaluate UniversalEPI’s sensitivity to genomic variation, specifically assessing whether UniversalEPI can detect allele-specific chromatin interactions driven by single-nucleotide variation, we tested the model on a well-characterized case involving SNP rs1800734. This variant was experimentally demonstrated to enhance *DCLK3* expression in colorectal cancer by strengthening interactions between a distal regulatory element encompassing rs1800734, the gene’s promoter, and its 3′ UTR^35^. Using ATAC-seq data from isogenic COLO320 cell lines homozygous for either the reference (G) or alternate (A) allele generated by Liu et al.^35^, we computed UniversalEPI’s Hi-C predictions. Consistent with experimental findings, UniversalEPI predicted stronger interactions between the SNP-containing region and both *DCLK3* promoter and 3′ UTR in the alternate allele background, as shown by maximum-confidence FC of predicted Hi-C values (Figure 2i). These results demonstrate that UniversalEPI can detect allele-specific chromatin interaction differences when paired with corresponding chromatin accessibility data.

### 2.3 Benchmarking UniversalEPI on unseen cell types using bulk inputs

We compared UniversalEPI against the leading method predicting raw chromatin interactions from bulk data, C.Origami^21^. While demonstrating high reported accuracy, C.Origami relies on additional CTCF ChIP-seq input^21^. To enable a fair comparison under limited-input settings, we additionally retrained C.Origami using only DNA sequence and ATAC-seq data. We also introduced three baselines that capture the effect of the distance between interacting elements and the variability between the train and test Hi-C interaction matrices (Methods). As the central goal of this work is to develop a model that generalizes to unseen cell types, we benchmarked the models in four cell lines not used for model training, using the same train-test split as above. Performance was evaluated based on the ability to predict experimental Hi-C interactions between accessible regions within 1 Mb, which was the maximum common receptive field across all existing methods.

We observed that UniversalEPI significantly outperformed the original version of C.Origami, which used CTCF ChIP-seq as an additional input, in predicting chromatin interactions in cell lines not seen during training (P < 0.05; Figure 3a, Figure S3). Performance declined further when C.Origami was retrained without CTCF input (P < 0.01; Figure 3a). UniversalEPI also outperformed the DNA sequence-only method Akita^14^, which is inherently limited in its ability to generalize to cell types not seen during training, and EPCOT^22^, which predicts observed-over-expected Hi-C using DNA sequence and chromatin accessibility (Figure S4). Notably, UniversalEPI was the only method to achieve Spearman’s correlations above 0.90 between predicted and experimental Hi-C values across all test cell lines (Figure 3b). On IMR90 and K562, UniversalEPI achieved comparable results to C.Origami, and it substantially outperformed C.Origami on HepG2 and GM12878 cell lines.

**Figure 3.**
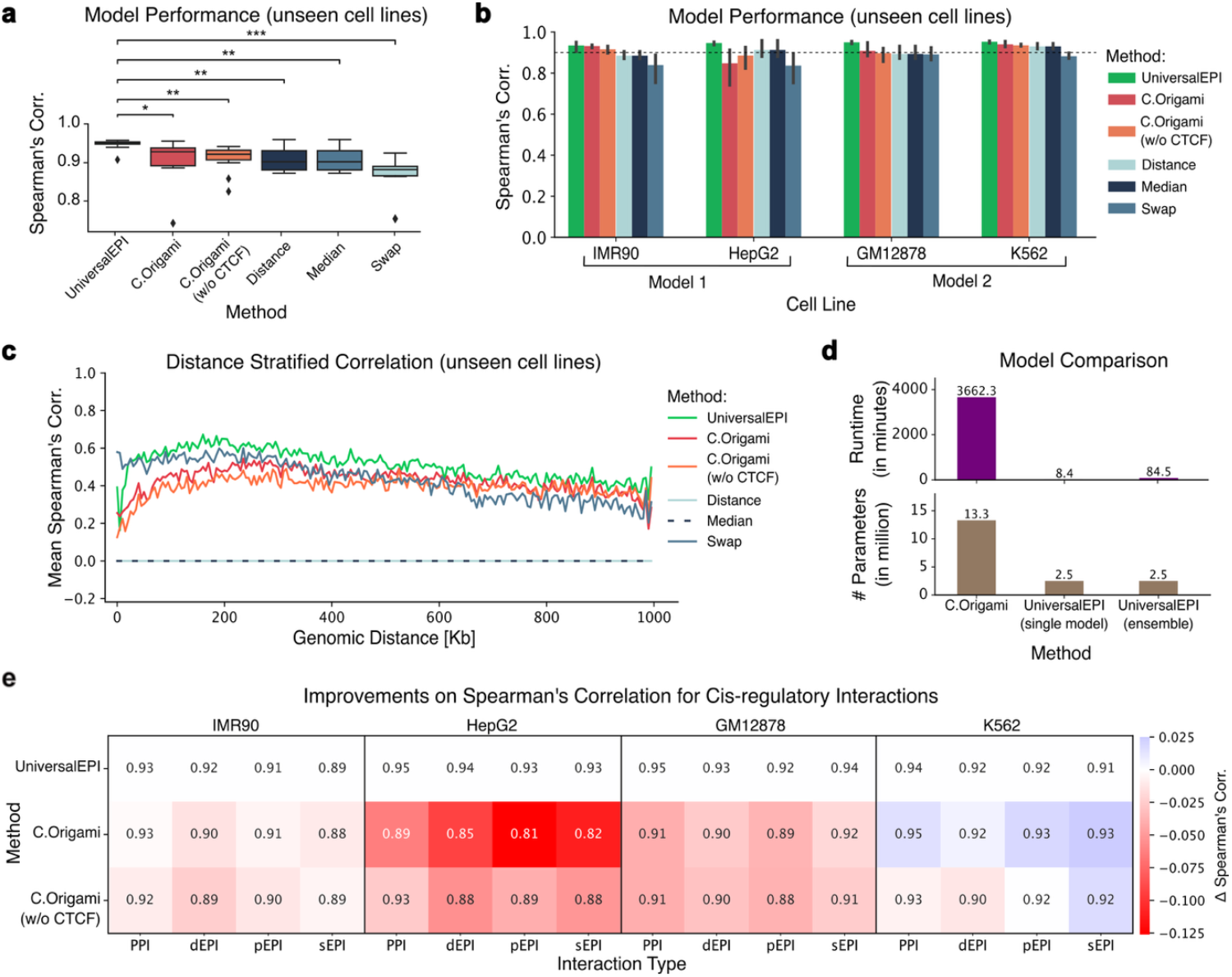
Benchmarking of the UniversalEPI method. **a,** Aggregated model performances for prediction of chromatin interactions in unseen cell types. Each point corresponds to Spearman’s correlation coefficient between predictions and ground-truth Hi-C for an unseen chromosome in unseen cell lines, IMR90 and HepG2 (Model 1), or GM12878 and K562 (Model 2). For each method, Model 1 is trained on GM12878 and K562 cells, whereas Model 2 is trained on IMR90 and HepG2 cells. A Wilcoxon signed rank test is used to compare the methods; ***: *P <* 0.001, **: *P <* 0.01, *: *P <* 0.05. **b,** Model performances for prediction of chromatin interactions in unseen cell types stratified by test cell line. The dotted line indicates Spearman’s correlation of 0.9. **c,** Line plot showing the mean distance-stratified Spearman’s correlation across four cell lines. **d,** Inference runtime and total number of model parameters are compared between different methods. The inference runtime is calculated for making the predictions using the test chromosomes of HepG2 cell line. **e,** Method performances for prediction of the Hi-C signal for different types of cis-regulatory interactions. The change in Spearman’s correlation is calculated with respect to UniversalEPI. Positive values suggest that C.Origami or its variant have higher scores than UniversalEPI (red). dEPI: distal Enhancer-Promoter Interactions (further than 2Kb), pEPI: proximal Enhancer-Promoter Interactions (within 2Kb), PPI: Promoter-Promoter Interaction, sEPI: Super Enhancer-Promoter Interactions.

Since the distance between the interacting elements is a major factor in determining the strength of interaction, we also compared the existing methods after removing the distance effect. This was done by calculating the distance-stratified correlation (Figure 3c), which assessed the ability of tested methods to capture biological features across several distance ranges. We observed that UniversalEPI predicted the strength of chromatin interactions at a given distance better than all the existing methods, including C.Origami. The possible reason for the lower performance of C.Origami could be overfitting on cell lines used in training. This was also confirmed by a higher performance of C.Origami on the test chromosomes of training cell lines as compared to UniversalEPI (Figure S5). In addition to a better performance than existing methods on cell types unseen during training, UniversalEPI offers an efficient solution with a small model size and a short inference runtime, making our method a more scalable alternative for large-scale applications (Figure 3d).

We compared UniversalEPI and C.Origami (with and without CTCF ChIP-seq as input) on their ability to predict specific types of EPIs and promoter-promoter interactions (PPIs) in the same test cell lines as above using experimental Hi-C as ground-truth. To study the ability of the models to capture distance-dependent interactions, we split EPIs into two categories – proximal EPI (pEPI, 200bp–2Kb) and distal EPI (dEPI, *>*2Kb). The annotations for pEPIs, dEPIs, and PPIs were obtained from the ENCODE library^36^. We also compared the two methods based on their ability to predict interactions between promoters and super-enhancers (sEPIs), which are clusters of active enhancers that often regulate cell identity genes^37^ and are defined for each cell line in the dbSuper database^38^. UniversalEPI outperformed both versions of C.Origami on all but the K562 cell line, where C.Origami with additional CTCF binding information showed slightly higher prediction accuracy than UniversalEPI (Figure 3e). Overall, we conclude that UniversalEPI generally captures important cis-regulatory interactions more accurately than C.Origami and that DNA sequence and bulk ATAC-seq data are sufficient to accurately predict cell-type-specific chromatin interactions.

Given UniversalEPI’s strong performance in predicting cell-type-specific cis-regulatory interactions using only DNA sequence and bulk ATAC-seq data, we deposited its log-transformed ICE-normalized predictions, derived from high-quality ATAC-seq data for 157 ENCODE cell lines and primary cells, to the UCSC Genome Browser (https://boevalab.inf.ethz.ch/resources/universalepi_pred_encode/ucsc_track_hub/). In addition to the raw predictions, we also provide z-score normalized Hi-C outputs to highlight long-range interactions. These publicly available tracks enable researchers to explore and validate regulatory interactions, supporting investigations into cell-type-specific chromatin architecture and gene regulation.

### 2.4 UniversalEPI captures chromatin dynamics during macrophage activation

UniversalEPI can be used to predict changes in regulatory chromatin interactions upon cell reprogramming. To demonstrate this, we analyzed the model’s predictions of chromatin interactions during activation of human macrophages in vitro. ATAC-seq and ground-truth Hi-C data were collected at eight time points following macrophage activation with lipopolysaccharide (LPS) and interferon-γ (IFN_*γ*_)^29^ (Figure 4a). In a zero-shot setting, Model 1 of UniversalEPI, which was trained on GM12878 and K562 cell lines, demonstrated high prediction accuracy, with Pearson’s and Spearman’s correlation with ground-truth Hi-C values exceeding 90% (Figure 4b).

**Figure 4.**
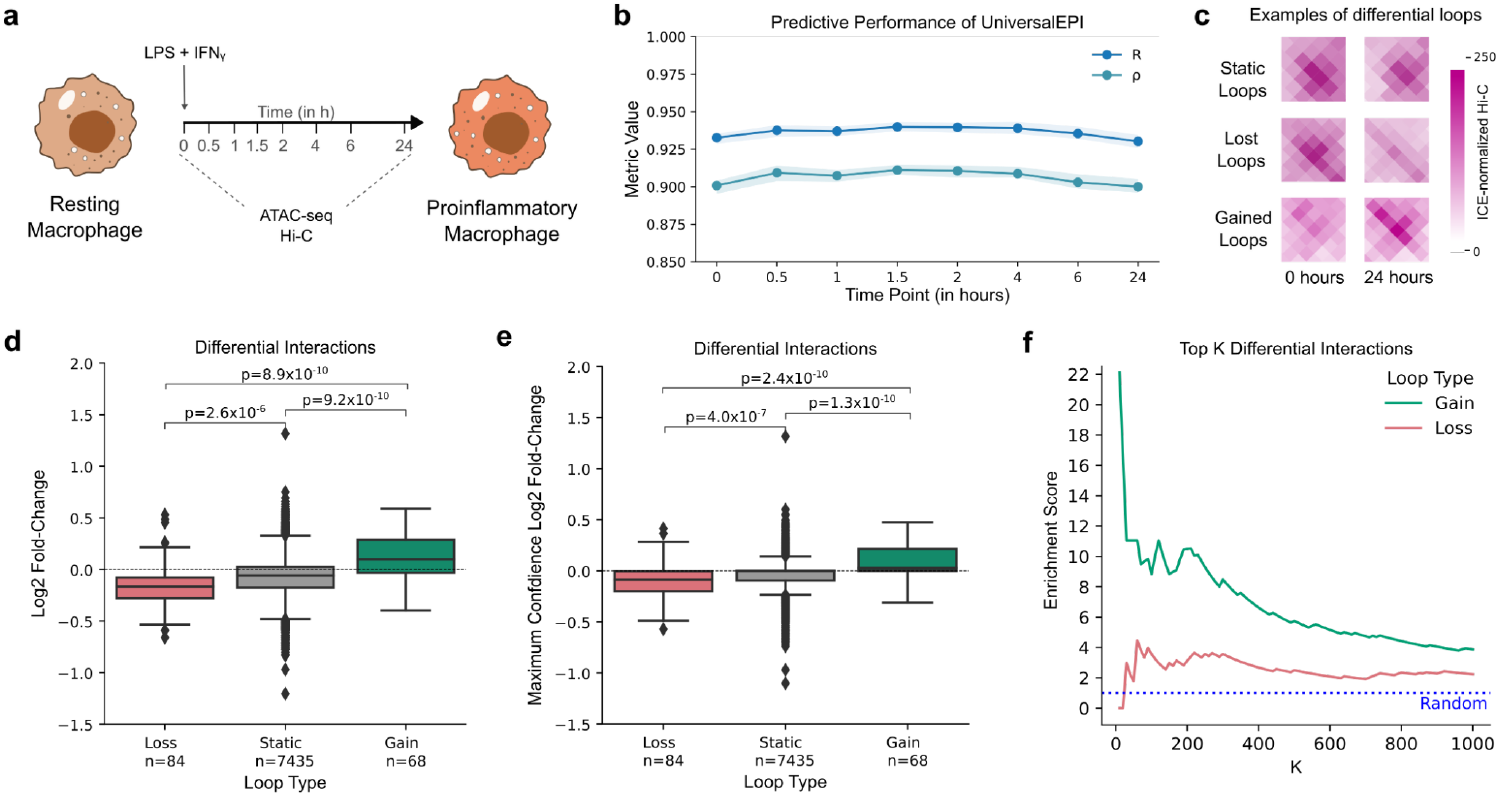
UniversalEPI captures chromatin dynamics during cell reprogramming. **a,** Experimental layout defined in Reed *et al*.^29^. Macrophages derived from the human THP-1 monocytic cell line are activated by treating with LPS + IFN_*γ*_in vitro and ATAC-seq and Hi-C were subsequently measured at 8 time points. **b,** Performance of UniversalEPI ensembles (trained on GM12878 and K562 cells) using ATAC-seq for each of the 8 time points, quantified by Pearson’s (*R*) and Spearman’s (*ρ*) correlation. **c,** Examples of static, lost, and gained loops in the chromatin between 0 and 24 hours identified by Reed *et al*. **d,** Differential interactions predicted by the UniversalEPI model between the 24-hour time point and the 0-hour for the differential loops. The significance is measured using a Mann-Whitney-Wilcoxon test with Bonferroni correction. **e,** The maximum-confidence log2 fold-change between Hi-C values measured by UniversalEPI ensemble predictions for the differential loops. The significance is measured using a Mann-Whitney-Wilcoxon test with Bonferroni correction. **f,** Gain and loss enrichment score over random for selecting a gained or lost loop, respectively, in the K most differential interactions as predicted by the UniversalEPI model. The blue dotted line represents the enrichment score of labeling a random interaction as gained or lost.

Reed *et al.* identified specific chromatin loops that exhibited differential interactions upon macrophage activation—lost, gained, or remaining static (Figure 4c). For these selected chromatin loops, UniversalEPI effectively identified positive and negative changes in the Hi-C signal during macrophage activation at 24h and 0h time points solely based on the variation in ATAC-seq inputs (Figure 4d). The application of the maximum-confidence log2 FC, based on the estimated prediction uncertainty, further improved the significance of predicted differential interactions (Figure 4e, Methods). Moreover, using the maximum-confidence log2 fold-change, UniversalEPI accurately distinguished gained and lost loops from static loops with a true positive rate that is at least four times higher than a random classification (Figure 4f). Therefore, we conclude that the unique feature of UniversalEPI – estimation of the maximum-confidence FC between conditions based on the reported uncertainty values – allows for accurate prediction of differential EPIs across conditions.

### 2.5 UniversalEPI predicts differential chromatin interactions using single-cell ATAC-seq data

To demonstrate the applicability to single-cell studies, we used UniversalEPI to predict chromatin structure using as inputs pseudo-bulks of single-cell ATAC-seq (scATAC-seq) data (Figure 5a). We compared the performance of UniversalEPI with ChromaFold^23^, the only other method that predicts bulk Hi-C from scATAC-seq and DNA sequence as inputs. Because ChromaFold was specifically designed to work on single-cell inputs, we did not include it in the above benchmarking on bulk ATAC-seq. scATAC-seq and Hi-C data from GM12878, K562, HepG2, and IMR90 cell lines were used in this analysis, with GM12878 and HepG2 cells employed for training both models. Keeping the experimental design similar to the ChromaFold study by Gao *et al.*^23^, we used Hi-C at 10Kb resolution with Hi-C-DC+ z-scores as targets^39^.

**Figure 5.**
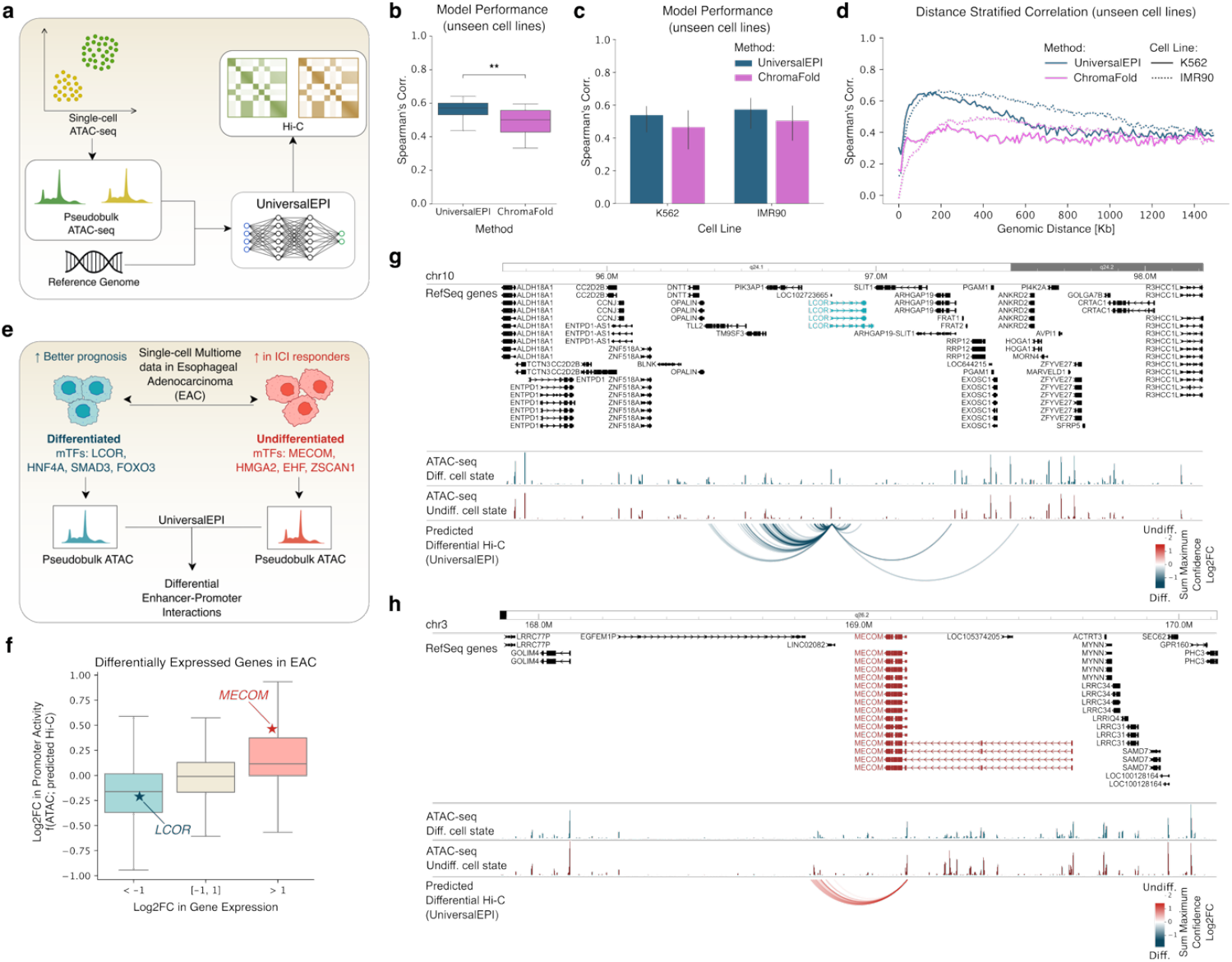
UniversalEPI reveals changes in chromatin structure in complex tissues using pseudo-bulk single-cell ATAC-seq data. **a,** UniversalEPI can predict cell-type-specific Hi-C-derived chromatin interactions using scATAC-seq data. **b,** Comparison between UniversalEPI and ChromaFold on their ability to predict bulk Hi-C from scATAC-seq using GM12878 and HepG2 cell lines as training. Spearman’s correlation is calculated on unseen chromosomes (chr5, chr18, chr20, chr21) of unseen cell lines (K562, IMR90). Wilcoxon signed-rank test is used to test significance. **: P ≤ 0.01 **c,** Performance comparison on unseen chromosomes stratified by unseen cell lines. **d,** Distance-stratified Spearman’s correlation based on the unseen chromosomes and cell lines. **e,** The 10x multiome data (scRNA-seq and scATAC-seq) of differentiated and undifferentiated malignant cells are obtained from human esophageal adenocarcinoma (EAC) tumors^40^. Master transcription factors (mTFs) for each program are identified in Yates *et al.*^40^. The scATAC-seq data from the candidate cells expressing each program are merged from multiple patient tumors to obtain transcriptional program-specific pseudobulk ATAC-seq profiles. These are then used as inputs to the pretrained UniversalEPI model to obtain Hi-C interactions for each program. **f,** A positive correlation is observed between the log2 fold-change (log2FC) in bulkified gene expression profiles and log2FC in estimated promoter activity. The blue star highlights the *LCOR* gene, which is a mTF for differentiated cells, whereas the red star highlights the *MECOM* gene, which is a mTF for undifferentiated cells. **g-h,** The change in predicted Hi-C interactions between accessible regions, measured by the sum of log2FC between undifferentiated and differentiated cells, are shown for the *LCOR* and *MECOM* gene promoters.

The two methods were compared based on their ability to predict Hi-C values on unseen chromosomes in the unseen cell lines IMR90 and K562. UniversalEPI significantly outperformed ChromaFold on both unseen cell lines (Figure 5b-c). UniversalEPI also obtained a higher distance-stratified correlation, especially for short-to-medium-range chromatin interactions (50-600Kb), highlighting the ability of our model to maintain its generalization capacity while extending to pseudo-bulk scATAC-seq data (Figure 5d).

Further, we used 10x multiome data (scATAC-seq and scRNA-seq) from 8 human esophageal adenocarcinomas (EACs)^40^ to assess potential differences in chromatin structure between two cancer cell states: differentiated and undifferentiated malignant cells expressing cNMF4 and cNMF5 transcriptional programs, respectively, as described in Yates *et al.*^40^. Pseudo-bulk ATAC-seq profiles were obtained from merging scATAC-seq data from all cells across patients expressing a given transcriptional program. These pseudo-bulk ATAC-seq profiles were then used to predict Hi-C interactions between accessible regions using the UniversalEPI model 1, pre-trained on GM12878 and K562 cell lines (Figure 5e).

Based on predicted Hi-C data, we defined values of promoter activity for each gene and compared them with measured gene expression from pseudobulk scRNA-seq. Promoter activity acts as a representative for gene expression and was defined as a function of ATAC-seq and the predicted Hi-

C. We modeled promoter activity as a combination of ATAC-seq promoter signal, ATAC-seq enhancer signals, and the predicted Hi-C interaction strengths between the promoter and enhancers (Methods). The derived promoter activity not only showed a high correlation with gene expression (Figure S6) but also a positive correlation with differential gene expression between undifferentiated and differentiated cells in EAC (Pearson’s *R* = 0.224, Figure 5f). Thus, UniversalEPI-derived differential chromatin interactions can indicate the strength of regulatory effects on gene expression. Yates *et al.* also identified master transcription factors (mTFs) for each program, including *LCOR* and *MECOM* in differentiated and undifferentiated EAC cells, respectively^40^. These TFs play a pivotal role in defining and maintaining malignant cell states. Specifically, *LCOR* expression drives tumor cell differentiation and reduces tumor growth, whereas low *LCOR* expression is associated with suppression of antigen processing and presentation, enabling resistance to immune checkpoint blockade (ICB) therapy in breast cancer^41,42^. Conversely, overexpression of *MECOM* has been linked with activating oncogene expression^43^, promoting stem cell-like properties, and reducing apoptosis in different cancers^44,45^. The UniversalEPI-predicted Hi-C signal for the two malignant cell states of EAC was in agreement with these findings, showing higher estimated promoter activity of *LCOR* in differentiated cells relative to the undifferentiated cells, in line with the increased *LCOR* gene expression, and vice versa for *MECOM* (Figure 5f).

Furthermore, using UniversalEPI-estimated uncertainty values, we identified significant differences in Hi-C interactions for the *LCOR* and *MECOM* promoters between the two malignant transcriptional states. In differentiated EAC cells, the *LCOR* promoter exhibited stronger interactions with proximal and distal enhancers both upstream and downstream of the gene (Figure 5g). Conversely, in undifferentiated EAC cells, the *MECOM* gene promoter formed strong interactions with downstream enhancers in its vicinity, presumably contributing to gene activation (Figure 5h). The uncertainty estimates allowed the model to discard low-confidence interactions, highlighting key enhancers responsible for gene regulation (Figure S7). This example illustrates the unique feature of UniversalEPI, which enables accurate identification of differential EPIs across conditions.

## 3 Discussion

In this work, we present UniversalEPI, an attention-based deep ensemble model that is trained on chromatin accessibility (ATAC-seq), DNA sequence, and TF binding data (ChIP-seq) from multiple cell lines to accurately predict chromatin interactions on unseen cell types without retraining solely based on DNA sequence and chromatin accessibility. Other models require several data modalities as inputs and, because of their large size, can only be trained on one or two cell types. UniversalEPI overcomes these drawbacks by operating on DNA sequence within open chromatin. By ignoring information in closed chromatin, other than assessing the distance between open chromatin regions, UniversalEPI is lightweight and demonstrates notable scalability (Figure 1). In contrast to other techniques, such as C.Origami, which was originally trained on a single cell line^21^, this enables our model to be trained on several cell lines concurrently in less than a day on a standard GPU and to have vastly improved inference time (Figure 3d). The latter will allow our model to be used on large datasets such as The Cancer Genome Atlas (TCGA) in the future, *e.g*., to assess the effects of non-coding variants on EPIs.

Using CNN layers and transformer blocks, UniversalEPI effectively captures the essential transcription factor binding motifs and spatial dependencies underlying interactions between regions of open chromatin. Thus, UniversalEPI consistently outperforms the state-of-the-art methods, including C.Origami, and shows Spearman’s correlation between the predicted and ground-truth Hi-C values on unseen cell types above 0.9 in all experimental settings (Figure 3b, Figure 4b). Furthermore, the model’s high prediction accuracy, demonstrated by the strong distance-stratified correlation between the predicted and experimental Hi-C values, confirms its sensitivity to biologically relevant information, which is critical for accurate modeling of chromatin architecture (Figure 2b, Figure 3c, Figure 5c).

By applying UniversalEPI to predict changes in chromatin architecture in human macrophages stimulated with LPS and IFN_*γ*_, we demonstrated it can perform in a zero-shot setting to find differential EPIs. Thus, UniversalEPI functions as a generalizable model that can identify modified chromatin interactions across conditions in cell types that are unseen during training (Figure 4). The model’s ability to detect dynamic changes in chromatin organization, which can be crucial in examining how different cell types respond to drugs and environmental stimuli, was demonstrated by its accurate identification of differential chromatin loops. UniversalEPI also displayed high performance against the only other model, ChromaFold, which was designed to predict bulk Hi-C using pseudo-bulk single-cell ATAC-seq data (Figure 5b-c). Furthermore, different chromatin looping patterns were identified by UniversalEPI for master transcription factor genes in EAC cells in different states of differentiation (Figure 5e-g). This feature means that UniversalEPI can be used to accurately investigate the characteristics and drivers of heterogeneity in chromatin organization that occurs in complex tissues and cancer cell states.

Its predictive power will enable UniversalEPI to be used to study changes in chromatin organization in the presence of non-coding mutations and structural variations, thereby helping to decipher the regulatory mechanisms of genetic diseases. In this work, we performed in silico inversions of CTCF binding motifs to reveal changes in chromatin interactions using UniversalEPI; future work could apply the same principles to large cancer datasets such as TCGA, which comprises data on chromatin accessibility and DNA sequence variation.

As UniversalEPI is trained to predict the Hi-C signal, the strong EPIs predicted by UniversalEPI are not strictly functional. It is known that enhancers and promoters can gain chromatin accessibility and interact even before gene expression is activated^10^. Additional data, such as profiling of H3K27ac or eRNA transcription, may be needed to assess the functionality of predicted strong EPIs.

Finally, the computational architecture of UniversalEPI sheds light on the ubiquitousness of chromatin folding mechanisms between promoters and enhancers in the cell types used for validation (lung fibroblasts, B lymphocytes, macrophages, chronic myelogenous leukemia cells, and hepatocellular carcinoma cells): the method demonstrates top performance while discarding all information about non-accessible chromatin and cell-type-specific TF binding motifs. Additional validation experiments may be required to check whether the model can show equally high performance in all cell types and whether highly specialized, cell-type-specific mechanisms that regulate chromatin interactions exist. As the numbers of ATAC-seq and Hi-C datasets increase, UniversalEPI can be used to explore these questions further.

In the future, UniversalEPI could benefit from being trained on more datasets to further improve its generalizability. Moreover, the first stage of UniversalEPI could be enhanced by incorporating techniques such as adversarial training^46–48^, which would additionally help prevent the model from capturing cell-type-specific information and improve its ability to generalize across cell types. Nonetheless, UniversalEPI achieves state-of-the-art performance in predicting EPIs in unseen cell types from only DNA sequence and chromatin accessibility profiles, and because of its unique feature of estimating prediction uncertainty, significant differential interactions can be detected across conditions. Therefore, UniversalEPI brings a major advancement in being able to predict and experiment on chromatin interactions in silico that can be widely applied to study complex regulatory landscapes across the genome.

## Methods

### Hi-C data processing

The raw hg38-aligned Hi-C files for four cell lines (GM12878, K562, IMR90, and HepG2) were obtained directly from the 4D Nucleome Data Portal^49^ under the accession codes listed in Supplementary Table 1. We then extracted the 5Kb resolution contact matrices. To remove any systematic biases, we applied ICE normalization^50^ using HiCExplorer v2.2.1^51^ with filter thresholds of −1.1 and 4.5. Since we are only interested in intra-chromosomal contacts, the ICE normalization was also done independently on each chromosome of each cell line.

The Hi-C contact matrices must be normalized across cell lines for the model predictions to be comparable to the training data. We achieved this by applying a distance-stratified robust z-score normalization using the healthy GM12878 cell line as a reference. GM12878 was selected as a reference cell line due to the high sequencing depth of its Hi-C matrix and hence, better data quality (approximately 759M interactions as compared to approximately 275M interactions in the other cell lines). For each chromosome of the GM12878 cell line, we stored the median and median absolute deviation (MAD) of all measured Hi-C interactions corresponding to regions that are *d* bins apart where *d* ∈ [0,800] (each bin being 5Kb of the genome). In some chromosomes, there were too few long-range interactions for accurate calculations of median and MAD. Hence, we fitted a B-spline, using SciPy’s splrep function^52^, setting the smoothing condition *s* to 5. This spline was then used to smooth our median and MAD curves across the genomic distance. We denote the smoothed median and MAD as 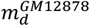 and 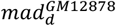 respectively. For a new cell type, e.g., the K562 cell line, the normalized

Hi-C score (ŷ) between regulatory elements in bins *i* and *j* is then calculated as done in equation (1),

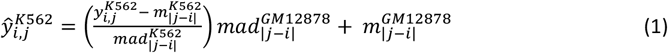

where *y* is the ICE-normalized Hi-C value. Figure S8 depicts the effect of this normalization on all cell lines.

Fudenberg *et al*^14^ applied Gaussian smoothing with unit variance to fill in the missing interactions and reduce the noise of the Hi-C experiment. However, this can lead to a significant reduction in strong Hi-C contacts in the presence of missing values at shorter distances. To mitigate this issue, we applied Gaussian smoothing with distance-dependent variance, which increased with distance (Figure S9). Particularly, we used four kernels with variances of 0.5, 0.65, 0.8, and 1 for interactions between regions that are closer than 25Kb, between 25Kb and 100Kb, between 100Kb and 250Kb, and greater than 250Kb respectively. Finally, we applied logarithmic transformation to the interaction scores. Several methods like Akita^14^ and ChromaFold^23^ also remove the effect of distance from the model by applying z-scores. While this approach highlights long-range interactions, we aim to capture the true biological interactions and hence, retain the effect of distance in our Hi-C matrices.

### ATAC-seq data processing

In this work, we used the GRCh38/hg38 human reference genome. ATAC-seq data from five human cell lines was used in this study consisting of healthy (GM12878 and IMR90) and cancer (K562, A549, and HepG2) cells. The carefully processed signal p-value bigwig profiles and the pseudoreplicated peak files were downloaded directly from the ENCODE portal (http://www.encodeproject.org/)”) under the accession codes listed in Supplementary Table 1. For multiple ATAC-seq peaks within the 500bp genomic region, we retain the peak with the highest read count and remove the remaining peaks. This resulted in approximately 175K peaks for each cell line.

To ensure consistency between the ATAC-seq bigwig profiles of different cell types, we normalized the bigwig signal using GM12878 as the reference cell line. First, we identified a conserved set of CTCF sites by selecting the common peaks from CTCF ChIP-seq tracks of all five cell lines. For each identified peak, we then obtained the signal values from the.bigwig files of each cell line using deepTools multiBigwigSummary v3.5.3^53^. Finally, we applied the Trimmed Mean of M-values (TMM) normalization of EdgeR^54^ using the extracted ATAC-seq signal values on these conserved CTCF sites to obtain the scaling factor for each cell line. GM12878 cell line was used as a reference while TMM normalization was applied. The normalized bigwig was then constructed by multiplying the scaling factor with the original signal track. The effect of normalization for all the cell lines can be observed in Figure S10.

### ChIP-seq data processing

The “narrow peak” files of each TF from the list of target TFs (CTCF, YY1, and SP1) for all five cell lines (GM12878, K562, IMR90, HepG2, and A549) were obtained from the ENCODE portal. The accession codes can be found in Supplementary Table 1. To account for the role of CTCF orientation in the formation and stability of TADs, we split CTCF ChIP-seq peaks based on their orientation. Specifically, we utilized the consensus motif MA0139.1 from the JASPAR database^55^ to derive a position-specific scoring matrix, which was subsequently convolved over the forward and reverse strands of the reference genome. To determine the optimal cutoff for the resulting convolved signal, we employed a false negative rate of 1%, based on empirical estimates^56^ of the intergenic nucleotide frequency background distribution. Signals larger than the cutoff indicated the presence of the consensus motif on the respective strand. Since the resolution of the ChIP-seq signal was lower than the motif length, a peak was labeled as *forward* and *backward* facing if both signals exceeded the cutoff value (18% of all peaks). Overall, 49% of the CTCF peaks were assigned to a motif on the forward strand and 48% to a motif on the reverse strand. For 20% of the peaks, no corresponding strand could be determined. Since these peaks were also highly correlated with low enrichment, they were dropped from the dataset during training.

### Model architecture

The architecture of UniversalEPI only accounts for the information coming from accessible chromatin regions (ATAC-seq peaks); all information about genomic regions between accessible regions is provided to the model via the value of the distance between ATAC-seq peaks. The model, therefore, predicts the entries in the interaction matrices only for the pairwise interactions between genomic regions containing ATAC-seq peaks. We propose a two-stage deep learning model to learn the mapping F: 𝕏 ↦ 𝕐, where 𝕏 is the space of genomic information about ATAC-seq peaks and 𝕐 is the space of interactions. Namely, for each *X* ∈ 𝕏, *X* = {*x*_1_,*x*_2_,..*,x*_401_} with *x*_*i*_∈ ℝ^1000×5^ being stacked one-hot encoded DNA sequences and ATAC-seq signal of one ATAC-seq peak, the mapping should produce *Y* = {*y*^(*i,j*)^ | *y*^(*i,j*)^ ∈ ℝ, *i* ≤ *j, i,j* = 101,*…,*300} ∈ 𝕐 with elements representing the interaction between the pair of peak locations *i* and *j* in the central 200 ATAC-seq peaks. Note that the entry *y*^(*i,j*)^, which represents the interaction between the pair of peaks locations *i* and *j*, depends not only on the data at those locations but also on the information across other peaks in their neighborhood.

The first stage learns genomic features for the peak regions, relevant to predicting generalized chromatin features across cell types, whereas the second stage predicts Hi-C values by modeling the interactions between the learned features at peak locations. A schematic overview of the proposed method is depicted in Figure S11. Below, we describe the two stages in detail.

#### Stage 1: Representation Learning

In the first stage, we train a representation network *f*_*θ*_(*x*), where *x* ∈ ℝ ^1000×5^, to predict binding affinity of the target TFs 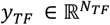 using the data across different cell lines, where *N*_TF_ is the number of the target TFs. This approach ensures that the learned features capture information that is both invariant across different cell lines and relevant for predicting the target Hi-C. Inspired by DeepC^13^, we adopt a convolution-based model for this purpose. The model consists of five convolutional layers with

{30,60,60,90,90} channels, and kernel sizes of {11,11,11,5,5} respectively. After each convolutional layer, max pooling with widths of {4,5,5,4,2} is applied to aggregate the learned features, which is followed by Leaky Rectified Linear Unit (LeakyReLU)^57^ with a slope of 0.2 to introduce non-linearity. A fully connected layer takes the output of the convolutional layers and maps to the target TFs. To prevent over-fitting, a 20% dropout is applied during training. The model is optimized by minimizing the empirical version of the mean squared error (MSE) defined as in equation (2),

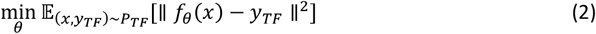

where paired *x* and *y*_TF_ are assumed to be sampled from a joint data distribution *P*_TF_.

The use of convolutional layers enables the network to effectively extract hierarchical features from the input data. After applying the non-linear activations, the outputs of convolutional blocks serve as intermediate representations and are used as the input for the second stage. We use a linear projection head for each feature layer to map it to the space of dimension *C*, which is set to 180. For one peak *x*, we obtain a set of *N*_*L*_projected features 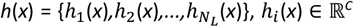, where *N*_*L*_ = 5 is the total number of convolutional blocks. Thus, for the subsequent 401 ATAC-seq peaks, we get the stacked projected features of the dimension 401 × *C* ×*N*_*L*_, which we denote by *h*(*X*). The feature selection module is then employed to select from *N*_*L*_features, as described in details below.

#### Stage 2: Hi-C Prediction

- *Automatic feature selection* is enabled by adopting the stochastic gating mechanism proposed by Yamada *et al*.^58^ to automatically select deep features learned during the first stage for predicting Hi-C interactions. Unlike traditional deterministic approaches, which rely on predefined criteria to select features, stochastic gating allows the model to explore a broader range of potential feature subsets and identify those crucial for the training task. Stochastic gating approximates the *ℓ*_0_ sparsity constraint by continuous relaxation of Bernoulli distribution. We jointly optimize the feature selection function *s*_*µ*_and the target network *g*_*ϕ*_, which predicts the Hi-C values. The total objective can be written as equation (3),

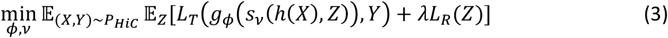

where *L*_*T*_and *L*_*R*_denote the target loss function and feature sparsity regularization, respectively, parameter *λ* controls the regularization strength, *Z* is a random sparsity-defining vector of size *N*_*L*_, *X* and *Y* are sampled from a joint data distribution *P*_HiC_. The components of *Z* provide the gating mechanism: For *k*-th feature *z*_*k*_= max(0,min(1,*ν*_*k*_+ *ϵ*_*k*_)), where *ν*_*k*_is learned and *ϵ*_*k*_ is sampled from 𝒩(0,*σ*^2^) with a predefined *σ* during training. Following Yamada *et al*.^58^, we set *σ* = 0.5. Herein we propose to use the stochastic gating *s*_*ν*_(*h*(*X*), *Z*) = {*s*_*ν*_(*h*(*x*_1_), *Z*), *s*_*ν*_(*h*(*x*_2_), *Z*), *…, s*_*ν*_(*h*(*x*_401_), *Z*)} at the feature level instead of the node level (the latter was initially proposed by Yamada *et al.*^58^) and aggregate the selected features using average pooling to reduce the model complexity. The gating mechanism is applied for each ATAC-seq peak 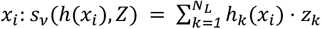. The sparsity regularization is the sum of the probabilities that the gates are active, which is equal to 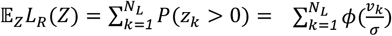, where *ϕ* is the Gaussian cumulative distribution function. During the inference stage, we set 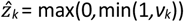.
- *Hi-C prediction network g*_*ϕ*_comprises a transformer-based encoder *g*_*ϕ*_^ENC^ that extracts information about the local and global context of the input sequence followed by a task-specific lightweight multilayer perceptron (MLP) decoder *g*_*ϕ*_^DEC^. The encoder consists of four multi-head attention blocks as proposed by Vaswani *et al.*^59^. We used four attention heads, a dropout rate of 0.1, and set the hidden dimension *d*_model_ equal to 32. To more accurately encode the genomic distance between input tokens, which is crucial for predicting interaction level, here we propose a genomic-distance-aware positional encoding. We compute the relative genomic distance between each token and a reference token and encode the distance using the sine-cosine positional encoding scheme^59^. Each token corresponds to a region centered around a single ATAC-seq peak. We designate the position of the highest ATAC-seq peak value as the token position. The middle token is chosen as the reference. We encode a maximum distance of 3 Mbp with a resolution of 500 bp. Note that the orientation is considered by assigning negative distances to the left peaks from the center and positive distances to the right peaks. We add a constant of 3 Mbp to all the distances to ensure they remain positive before the sine-cosine encoding. The encoder extracts a representation for each token, capturing its relation with the other tokens in the input sequence. The output of *g*_*ϕ*_^ENC^ is defined as *E*_*ϕ,ν*_ = {*e*_1_,*e*_2_,*…,e*_401_} with the *i*-th token representation 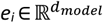. To compute the Hi-C interaction value between two peaks locations *i* and *j*, the encoder output of the *i*-th and *j*-th token are concatenated and fed into the decoder predicting Hi-C value, *i.e.*, 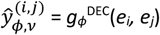 and *e*_*j*_depend on *ϕ,ν*. We use two MLP layers with ReLU activation for the decoder. During training, sequences are randomly flipped to ensure that the decoder remains invariant to the order of the tokens. The target loss *L*_*T*_between the true interactions *Y* and the estimated interactions Ŷ_*ϕ,ν*_in equation (3) is defined in equation (4),

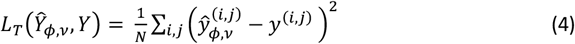

where *i,j* are encoded following the DeepC approach^13^ in a vertical, zigzag pole over the center of the sequence window (Figure 1c). *N* is the total number of interaction pairs. During training the empirical version of the loss (equation (3)) is optimized in both *ν* and *ϕ*.
- *Auxiliary information* can be optionally used for Hi-C prediction. Since the Hi-C values between two bins are also dependent on the mappability of the independent bins^60^, we also include the 36bp-mappability track, obtained from the UCSC Genome Browser^61,62^, as input to the Stage 2 of UniversalEPI. First, we use a linear head to embed auxiliary information. We set the embedding dimension the same as the hidden dimension of the transformer. The embedded vector is then concatenated with the input embeddings, resulting in a new input to the attention layers. This method allows the use of any auxiliary information, such as mappability and ATAC-seq.

### Uncertainty estimation

We incorporate both *aleatoric* and *epistemic* uncertainty estimation into UniversalEPI, providing information on the reliability of model predictions. To capture the *aleatoric* uncertainty, we assume that each observed log Hi-C value is a sample from a Gaussian distribution. Instead of the point estimate Ŷ_*ϕ,ν*_ comprised of 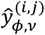 provided by the entire Hi-C prediction component *g*_*ϕ*_∘ *s*_*ν*_, we model the parameter of the distribution: the mean 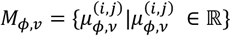 and the variance 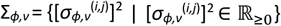 between two peaks locations *i* and *j*, which are structured the same way as *Y*. We apply an exponential activation function to the variance output to guarantee non-negativity. Instead of the MSE loss *L*_*T*_defined in equation (4), herein we minimize in *ϕ* and *µ* the negative log-likelihood loss weighted by the *β-*exponentiated variance estimates^27^, referred to as *β*−NLL loss, which is defined in equation (5),

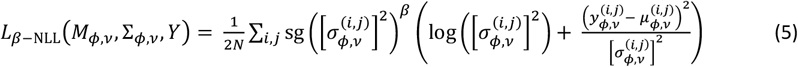

with the stop-gradient operator sg, and the parameter *β* controlling the trade-off between the regression accuracy and log-likelihood estimation. We set *β* = 0.5 following^27^, which has been empirically found to provide the best trade-off. To estimate *epistemic* uncertainty, we utilize the deep ensemble method^28^ by training *K* models with different random initializations, resulting in an ensemble {(*ϕ*_*k*_,*ν*_*k*_), *k* = 1,..*,K*}. The total predictive uncertainty 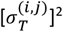 is a sum of aleatoric 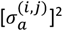 and epistemic 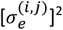 uncertainties. It is quantified by combining these components as described in equation (6),

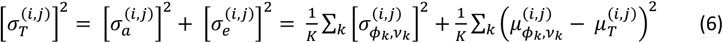

with the empirical estimate of the mean 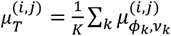. We use an ensemble size of *K* = 10 for the reported results.

### Training details

Our method is implemented with the PyTorch framework^63^. The training of the proposed method involves a two-stage model, where each stage was trained separately with different training setups. The TF prediction network was trained with Adam^64^ optimizer using a learning rate of 10^−4^, a weight decay of 10^−4^, with a batch size of 1024 over 100 epochs. The best model was identified using Pearson’s correlation score on the validation dataset. For each active binding site, we select a 1Kb segment of DNA sequence around the peak center and the corresponding ATAC-seq as input. Transcription factors CTCF, YY1, and SP1 were empirically chosen as target TFs based on their ability to assist in chromatin organization prediction (Figure S12). ZNF143 was initially considered another target TF candidate^65-67^. However, a recent study^68^ found the commonly used antibody of ZNF143 cross-reacting with CTCF, challenging reported associations between ZNF143 binding and chromatin looping. This, along with the fact that including ZNF143 did not improve Hi-C prediction accuracy (Figure S12), led us to exclude this TF from further consideration in our work. The first stage model was trained on the GM12878 and K562 cell lines. We randomly split the chromosomes into training, validation, and test sets. Chromosomes 5, 12, 13, and 21 were used for validation, 2, 6, and 19 for testing, and the remaining chromosomes for training. After training, the backbone of the TF prediction network was used as a feature extractor, and its model parameters were frozen.

In the second stage, the Hi-C prediction network was trained using the AdamW^69^ optimizer with a learning rate of 10^−3^. We set the feature sparsity regularization *λ* to 0.01, which was tuned to select the minimum number of feature layers necessary. Since distance between elements is highly predictive of the Hi-C values, we employed a position embedding upscaling by a factor of √*d*_*model*_. We trained two sets of models, one set uses GM12878 and K562 cell lines for training, and IMR90 and HepG2 cell lines for testing whereas the other uses HepG2 and IMR90 cells for training and GM12878 and K562 cells were held out for testing. The second stage model was trained using the same chromosome split as the first stage. We found that the Hi-C data at 5Kb resolution also contains arbitrary bins with very few reads mapped to them (Figure S13). To ensure that these noisy samples do not affect our analysis, we removed these and their one-hop neighboring bins from all datasets. To increase robustness, we augmented the accessible regions in the training dataset with 10% of inaccessible regions of length 1Kb each that were chosen randomly from the genome ensuring that the model can also predict accurately for various inaccessible regions, which can arise due to mutations. The combined regions were then used to form the 401 consecutive peaks which is the input of UniversalEPI. We trained the network for 20 epochs and chose the model for evaluation with the highest Spearman’s correlation on the validation data. Training the Hi-C prediction network took 12 hours on 0.2 of an A100 GPU with a batch size of 32, and 24 hours on a single RTX2080Ti with a batch size of 16.

### Evaluation details

UniversalEPI was evaluated on unseen chromosomes of unseen cell types. To ensure that reliable targets were used for evaluation, we first flagged the bins that overlap with unmappable (https://storage.googleapis.com/basenji_barnyard2/umap_k36_t10_l32_hg38.bed) and blacklisted (https://storage.googleapis.com/basenji_barnyard2/hg38.blacklist.rep.bed) regions as reported by Kelley *et al*^70^. We then removed the interactions containing a flagged endpoint. Finally, we calculated Spearman’s correlation and Pearson’s correlation using the smoothed log Hi-C of the remaining interactions.

To understand the biological information captured by our model, we removed the strong influence of distance on Hi-C prediction by calculating distance-stratified correlation. Specifically, we calculated the correlation using all interactions that lie at a particular distance *d* away from each other, where *d* lies between 0 and 2Mb with a step size of 5Kb.

### State-of-the-art methods

In this section, we describe the configurations of the state-of-the-art methods C.Origami^21^,Akita^14^ and EPCOT^22^, against which we compare our approach. We did not compare against DeepC^13^, as it has approximately 10 times more parameters than Akita and the two models are shown to perform comparably^14^.

Various existing methods follow different Hi-C preprocessing techniques, predict Hi-C up to different distances and at different resolutions. For example, Akita predicts observed over expected Hi-C at 2048bp resolution up to 1Mb, EPCOT also predicts observed over expected Hi-C at 5Kb resolution up to 1Mb, whereas C.Origami retains the effect of distances and predicts Hi-C at 10Kb resolution capturing interactions up to 2Mb away. Moreover, Akita employs 2D Gaussian smoothing which is not used by C.Origami.

To mitigate these differences, we re-trained C.Origami on our preprocessing of 5Kb Hi-C data and tuned its hyperparameters based on its performance on validation chromosomes of seen cell lines. We used the open-sourced code as a starting point. We did not modify the loss functions, optimizers, or learning rate schedulers. In case of C.Origami, we reduced the input size from 2Mb to 1Mb to compensate from increase in Hi-C resolution from 10Kb to 5Kb. This majorly led to changes in the hidden layer sizes and number of convolution layers in the encoder. To obtain the C.Origami model without the CTCF ChIP-seq as input, we simply excluded this track from the inputs without changing the configuration of the model. Finally, both versions of C.Origami models were trained for a maximum of 80 epochs. The top-performing model was selected based on the validation loss and used for inference.

Since Akita predicts the natural logarithm of observed over expected Hi-C corresponding to the input size of 1Mb, we first exponentiated the predictions and multiplied them with expected Hi-C before reapplying the natural logarithm operation. This converts Akita’s predictions to log raw Hi-C. We used the pre-trained Akita model to predict Hi-C interactions for K562 and HepG2 cell lines, as these cell lines were not included in its training. The average prediction across all training cell lines is used for both unseen cell lines. As Akita predicts the Hi-C contact map at 2048bp resolution, we apply adaptive average pooling to rescale the contact maps at 5Kb resolution, allowing us to make a fair comparison between UniversalEPI and Akita.

Finally, we used the pre-trained EPCOT models, one trained using the data from GM12878 cell line and the other using HFF cell line. Both models were trained using ATAC-seq and hg38 DNA sequence. Like Akita, EPCOT also predicts observed over expected Hi-C using 1Mb DNA sequence and chromatin accessibility as input. Therefore, we multiplied the model predictions with expected Hi-C to obtain the raw Hi-C signal. Since the sequence encoder of EPCOT is trained using data from GM12878, K562, and HepG2 cell lines, we compared the model predictions on the IMR90 cell line. Using IMR90’s ATAC-seq as input, the average prediction of the two pre-trained EPCOT models was considered as the final prediction.

### Baseline methods

We define 3 baseline methods, (1) Distance, (2) Median, and (3) Swap. These are computationally very simple as compared to the complex deep learning methods.

As the Hi-C interactions between two genomic segments heavily depend on the distance between the segments, we introduce the Distance baseline which only depends on the genomic distance between the open regions. In particular, we define the prediction between regions *I* and *j* in equation (7),

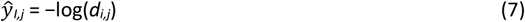

where *d*_*i,j*_is the genomic distance (in 5Kb bins) between regions *i* and *j*.

Similarly, we define the Median baseline where prediction is the median of all the interactions between the open regions in the training data that are at a particular distance away. Mathematically, the prediction for interaction between regions *i* and *j*, that are *d*_*i,j*_bins away, is given by equation (8),

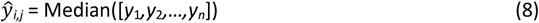

where *y*_1_ *…y*_*n*_are Hi-C interactions from the training data between all pairwise open regions that are *d*_*i,j*_bins apart.

Finally, we define the third baseline called Swap to capture the similarity in Hi-C across cell types. Here, we take the mean of Hi-C interactions from all the cell lines in the training data and use this as the predicted Hi-C matrix. If GM12878 and K562 are taken as training cell lines, then the predicted Hi-C interaction for the new unseen cell line would be given by equation (9).

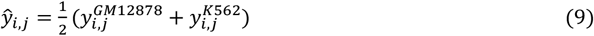

### TAD annotations

TADs were identified using boundary calls from the 4DN data portal^49,71^. We defined a consecutive pair of boundary calls (*i,j* where *i < j*) as a TAD if there was an overlapping ATAC-seq peak and a CTCF ChIP-seq peak within 10Kb of boundary calls *i* and *j*. Additionally, we required that the CTCF orientation is forward for *i* and reverse for *j*. The CTCF orientation was determined following the same approach used to prepare target data for the first stage of UniversalEPI. Applying these criteria yielded approximately 200 TADs per cell line.

### In-silico inversion of TAD boundary

For each identified TAD, we modified the ATAC-seq peak at boundary *j* by reversing the ATAC-seq signal and mappability profile and by taking the reverse complement of the DNA sequence, resulting in a modified peak. We then extracted features from this modified peak using the first stage of

UniversalEPI. These features, along with those from the 400 neighboring peaks, were used as input to the second stage of UniversalEPI.

Since interactions between two 5Kb bins can be influenced by multiple accessible chromatin regions within the two bins, it is challenging to isolate the effect of inverting a TAD endpoint in such cases. To address this, we focused our analysis on TADs with a unique ATAC-seq peak within 5Kb of each endpoint.

### Availability of Hi-C predictions on ENCODE

We generated the Hi-C predictions corresponding to all possible ATAC-seq data from the ENCODE portal. We included all human cell lines and primary cells that were not genetically perturbed and had sufficient read depth. All the datasets are summarized in Supplementary Table 2. This resulted in 116 ATAC-seq profiles for different cell lines and 41 ATAC-seq profiles for primary cells.

For each profile, the Hi-C predictions were made using the pretrained UniversalEPI models. The predictions were exported as *bigInteract* files with ensemble mean as interaction strength. The estimated uncertainty was reported in the interaction label. Along with the log-transformed Hi-C, we also generated the z-score Hi-C interactions as a separate track, highlighting the strong long-range interactions. For each predicted interaction *y*_*i,j*_ between regions *i* and *j*, we calculate the z-score normalized Hi-C by equation (10), where *μ*_*d*_ and *σ*_*d*_ are mean and standard deviation of all interactions that are at *d* = |*j* − *i*| distance away in the same chromosome.

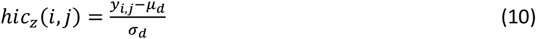

To ensure that only the strong interactions are viewed by the users, we set a threshold of 5 for ICE-normalized Hi-C predictions (or 1.8 after the log-transformation) and 0 for z-score normalized Hi-C predictions. A total of 314 tracks (157 log-transformed ice-normalized Hi-C + 157 z-score normalized Hi-C) were then submitted to UCSC Genome Browser as a public track hub enabling interactive visualization of predicted chromatin interactions in different cellular contexts. The URL to each track is also reported in Supplementary Table 2.

### Macrophage activation data processing

We validated UniversalEPI on the data collected for activated macrophages derived from the THP-1 monocytic cell line. Since we require the ATAC-seq peak calls and the signal p-value bigwig track, we used the raw ATAC-seq paired-end.fastq files provided by Reed *et al.*^29^ and performed preprocessing similar to the ones done by the authors. Adaptors and low-quality reads were first trimmed using Trim Galore! v0.6.10. Reads were then aligned using BWA mem. Samtools v1.13^72^ was then used to sort the aligned reads. Duplicated reads were removed using PicardTools whereas mitochondrial reads were filtered using Samtools. The replicates for each timestamp were merged using Samtools. Finally, ENCODE’s ATAC-seq pipeline^73^ was used to obtain the peaks and signal p-value bigwig track. Deduplication was done as described before for the generated peak files. This resulted in approximately 200K peaks for each time point. For the generated bigwig tracks, edgeR’s TMM normalization^54^ was done with the GM12878 cell line as the reference (Figure S10).

The.hic files, for each of the 8 different time stamps, were directly obtained from Reed *et al*^29^. Nearly 370M interactions were observed on average for each of the time points. ICE normalization^50^ was applied to each.hic file followed by the z-score normalization using the GM12878 cell line as the reference (as described above).

### Differential loop prediction

For the validation of the ability of UniversalEPI to identify differential loops, we used the selected loops provided by Reed *et al*. The *gain.early* and *gain.late* were merged to form the set of gained loops. A similar approach was used for lost loops. To ensure a reliable set of differential loops, the sets of gained, static, and lost loops were further filtered based on fold-change (*FC*) of ICE-normalized Hi-C between the 24 hours and 0 hour time points. Specifically, lost loops with *FC <* 0.67, gained loops with *FC >* 1.5, and static loops with *FC* ∈ [0.67,1.5] were retained.

The ATAC-seq peak sets at 0th and 24th hour time points were merged to get a set of regions for which Hi-C signal was predicted for the two time points using UniversalEPI. If multiple peaks existed within 500bp of each other, the peak with the maximum overall enrichment was retained resulting in a total of 235,690 peaks. UniversalEPI predictions were then independently calculated on this merged peak set using ATAC-seq bigwig signals for the two time points. Finally, the maximum-confidence log2 fold-change was calculated between the predictions at these two time points. Assuming UniversalEPI predicted an interaction for two time points *t*_1_ and *t*_2_ with means *µ*_1_ and *µ*_2_ and variances 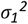 and 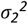 respectively, a parameter *d* was defined as 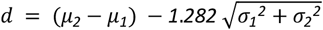 for *μ*_*2*_> *μ*_*1*_ and 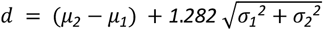 otherwise. Here, *d* quantifies whether the observed difference exceeds the expected variability, with 1.282 representing the z-score for a one-tailed 90% confidence level. The maximum-confidence log2 fold-change between time points *t*_2_ and *t*_1_ was set to 0 if *d* < *0* and *μ*_*2*_> *μ*_*1*_ or *d* > *0* and *μ*_*2*_≤ *μ*_*1*_, indicating the change was not statistically significant. In case of no overlap, the maximum-confidence log2 fold-change was given by d. This corresponds to log2 fold-change being at least d with 90% model confidence.

### Comparison with ChromaFold

To compare UniversalEPI’s performance against ChromaFold^23^, we trained and evaluated both models using the same input single-cell ATAC-seq (scATAC-seq) data. Since UniversalEPI is designed to work with bulk ATAC-seq signal and ATAC-seq peaks, we first bulkified the scATAC-seq data. The bam files were directly downloaded from ENCODE (Supplementary Table 1) and converted to signal and peak files using the techniques explained in the sections above. These bulkified ATAC-seq signals and peaks were used to train the UniversalEPI model and the evaluation of both models.

Both the models were trained using data from GM12878 and HepG2 cell lines and evaluated on K562 and IMR90 cell lines. We followed the same processing as introduced in Gao *et al.*^23^ *i.e.* we used ICE-normalized Hi-C at 10Kb resolution. Z-scores were then calculated using HiC-DC+^39^ and the resulting values were clipped between-16 and 16. Chromosomes 5, 18, 20, and 21 were used for testing whereas chromosomes 3 and 15 for validation, and the remaining chromosomes for training, as done in Gao *et al*^23^. This allowed us to directly use the preprocessing, model, and training scripts from the source code of ChromaFold.

The models were compared using interactions between accessible chromatin regions. The interactions were also filtered based on unmappable and blacklisted regions. UniversalEPI and ChromaFold were also evaluated based on distance-stratified correlation, evaluating the performance of these methods in predicting interactions between genomic regions located up to 1.5Mb away from each other.

### Esophageal adenocarcinoma data processing

The 10x multiome (single-nuclei ATAC-seq and single-nuclei RNA-seq) profiles of 10 esophageal adenocarcinoma (EAC) patients were obtained from Yates *et al*^40^. Using the cell scores from Yates *et al.*, we selected the top 20% unique cells in each of the differentiated (cNMF4) and undifferentiated (cNMF5) programs as the representative cells.

The representative cell barcodes are extracted from the individual patients’.bam file, and the resulting patient-specific.bam files are merged to obtain the pseudo-bulk ATAC-seq.bam. For each program, ATAC-seq bam files were then used to obtain the signal p-values bigwig and narrowpeak files using ENCODE’s ATAC-seq pipeline^73^ as done before. The ATAC-seq peaks were merged for the two programs to obtain a common set of peaks for downstream differential analysis. This was followed by deduplication (as done for datasets before) which resulted in a total of 280,368 peaks. The signal p-value bigwigs were normalized using edgeR’s TMM normalization^54^ was done with the GM12878 cell line as the reference (Figure S10).

The pseudo-bulk gene expression is obtained by summing the gene counts from all representative cells. Finally, the gene expression counts for each program were converted to counts-per-million (CPM) to mitigate the library-size bias.

### Promoter activity

The promoter activity (*A*_*P*_) was obtained using the accessibility of the promoter (ATAC_*P*_), accessibility of all the interacting enhancers (ATAC_*E*_), and the interaction strength between the enhancer and promoter (Hi-C_*E,P*_). The accessibility of each enhancer was extracted as the maximum signal in the vicinity (±50bp) of the gene transcriptional start site or the point of maxima called by peak calling of MACS2 for the enhancers. Specifically, we defined the promoter activity as a linear combination of the promoter accessibility and the Hi-C weighted by the total accessibility of the enhancer and promoter. Inspired by Loubiere *et al.*^74^, the total accessibility of enhancer and promoter was defined using the multiplicative model. This is mathematically given by equation (11),

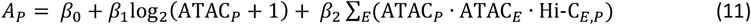

where *β*_0_ is the intercept and *β*_1_, and *β*_2_ are the coefficients, all of which are learned by a lasso regressor on randomly selected 80% of protein-coding genes in IMR90 cells. To mitigate the effect of weak interactions, only those enhancers were considered that had a predicted log Hi-C interaction of greater than 1 with the promoter.

## Data Availability

All the raw and processed datasets corresponding to the cell lines were obtained from publicly accessible databases including ENCODE (http://www.encodeproject.org/)”), 4DNucleome (4DN) (https://data.4dnucleome.org/), and Gene Expression Omnibus (GEO) (http://www.ncbi.nlm.nih.gov/geo/)”) with accession codes mentioned in Supplementary Sheet 1. This includes the THP-1 cell line used for macrophage activation generated in Reed *et al*. The raw single-cell ATAC-seq and single-cell RNA-seq datasets for cell differentiation in esophageal carcinoma were obtained from database of Genotypes and Phenotypes (dbGaP) (http://www.ncbi.nlm.nih.gov/gap/) with accession number phs003438.v1.

The precomputed Hi-C predictions for the 157 ENCODE ATAC-seq tracks of cell lines and primary cells are made available as a track hub for the UCSC Genome Browser at https://boevalab.inf.ethz.ch/resources/universalepi_pred_encode/ucsc_track_hub/hub.txt. The mapping between the generated interaction files and ENCODE datasets can be found in Supplementary Table 2.

## Code Availability

The code for UniversalEPI is made available at https://github.com/BoevaLab/UniversalEPI and deposited to https://doi.org/10.5281/zenodo.14622040. The tutorial is available at https://github.com/BoevaLab/UniversalEPI/wiki.

## Acknowledgements

This project is partially funded by the Swiss Data Science Center (SDSC) collaborative projects grant (C22-09); AG is funded by the Swiss Government Excellence Scholarship (ESKAS-Nr: 2021.0468).

## Competing interests

F.J.T. consults for Immunai Inc., Singularity Bio B.V., CytoReason Ltd and Omniscope Ltd, and has ownership interest in Dermagnostix GmbH and Cellarity. I.L.I. currently works at Bioptimus.

## Notes

### Summary of Updates

Added new results on allele-specific Hi-C prediction and created a resource of predicted Hi-C for 157 ENCODE cell lines and primary cells using UniversalEPI.

## References

1. Dekker, J., Marti-Renom, M. A. & Mirny, L. A. Exploring the three-dimensional organization of genomes: interpreting chromatin interaction data. Nat Rev Genet 14, 390–403 (2013).

2. Cavalheiro, G. R., Pollex, T. & Furlong, E. E. To loop or not to loop: what is the role of TADs in enhancer function and gene regulation? Current Opinion in Genetics & Development 67, 119–129 (2021).

3. Schoenfelder, S. & Fraser, P. Long-range enhancer–promoter contacts in gene expression control. Nat Rev Genet 20, 437–455 (2019).

4. de Wit, E. et al. CTCF Binding Polarity Determines Chromatin Looping. Mol Cell 60, 676–684 (2015).

5. Weintraub, A. S. et al. YY1 Is a Structural Regulator of Enhancer-Promoter Loops. Cell 171, 1573-1588.e28 (2017).

6. Deshane, J. et al. Sp1 regulates chromatin looping between an intronic enhancer and distal promoter of the human heme oxygenase-1 gene in renal cells. J Biol Chem 285, 16476–16486 (2010).

7. Yang, Y., Zhang, R., Singh, S. & Ma, J. Exploiting sequence-based features for predicting enhancer– promoter interactions. Bioinformatics 33, i252–i260 (2017).

8. Hsieh, T.-H. S. et al. Enhancer-promoter interactions and transcription are maintained upon acute loss of CTCF, cohesin, WAPL, and YY1. 2021.07.14.452365 Preprint at 10.1101/2021.07.14.452365 (2021).

9. Uyehara, C. M. & Apostolou, E. 3D enhancer-promoter interactions and multi-connected hubs: Organizational principles and functional roles. Cell Rep 42, 112068 (2023).

10. Pollex, T. et al. Enhancer-promoter interactions become more instructive in the transition from cell-fate specification to tissue differentiation. Nat Genet 56, 686–696 (2024).

11. Akdemir, K. C. et al. Disruption of chromatin folding domains by somatic genomic rearrangements in human cancer. Nat Genet 52, 294–305 (2020).

12. Feng, Y. & Pauklin, S. Revisiting 3D chromatin architecture in cancer development and progression. Nucleic Acids Research 48, 10632–10647 (2020).

13. Schwessinger, R. et al. DeepC: predicting 3D genome folding using megabase-scale transfer learning. Nat Methods 17, 1118–1124 (2020).

14. Fudenberg, G., Kelley, D. R. & Pollard, K. S. Predicting 3D genome folding from DNA sequence with Akita. Nat Methods 17, 1111–1117 (2020).

15. Zhou, J. Sequence-based modeling of three-dimensional genome architecture from kilobase to chromosome scale. Nat Genet 54, 725–734 (2022).

16. Li, W., Wong, W. H. & Jiang, R. DeepTACT: predicting 3D chromatin contacts via bootstrapping deep learning. Nucleic Acids Res 47, e60 (2019).

17. Zhang, S., Chasman, D., Knaack, S. & Roy, S. In silico prediction of high-resolution Hi-C interaction matrices. Nat Commun 10, 5449 (2019).

18. Yang, R. et al. Epiphany: predicting Hi-C contact maps from 1D epigenomic signals. Genome Biol 24, 134 (2023).

19. Whalen, S., Truty, R. M. & Pollard, K. S. Enhancer-promoter interactions are encoded by complex genomic signatures on looping chromatin. Nat Genet 48, 488–496 (2016).

20. Chen, K., Zhao, H. & Yang, Y. Capturing large genomic contexts for accurately predicting enhancer-promoter interactions. Brief Bioinform 23, bbab577 (2022).

21. Tan, J. et al. Cell-type-specific prediction of 3D chromatin organization enables high-throughput in silico genetic screening. Nat Biotechnol 41, 1140–1150 (2023).

22. Zhang, Z., Feng, F., Qiu, Y. & Liu, J. A generalizable framework to comprehensively predict epigenome, chromatin organization, and transcriptome. Nucleic Acids Research 51, 5931–5947 (2023).

23. Gao, V. R. et al. ChromaFold predicts the 3D contact map from single-cell chromatin accessibility. Nat Commun 15, 9432 (2024).

24. Ong, C.-T. & Corces, V. G. CTCF: an architectural protein bridging genome topology and function. Nat Rev Genet 15, 234–246 (2014).

25. Makhlouf, M. et al. A prominent and conserved role for YY1 in Xist transcriptional activation. Nat Commun 5, 4878 (2014).

26. O’Connor, L., Gilmour, J. & Bonifer, C. The Role of the Ubiquitously Expressed Transcription Factor Sp1 in Tissue-specific Transcriptional Regulation and in Disease. Yale J Biol Med 89, 513–525 (2016).

27. Seitzer, M., Tavakoli, A., Antic, D. & Martius, G. On the Pitfalls of Heteroscedastic Uncertainty Estimation with Probabilistic Neural Networks. Preprint at 10.48550/arXiv.2203.09168(2022).

28. Lakshminarayanan, B., Pritzel, A. & Blundell, C. Simple and Scalable Predictive Uncertainty Estimation using Deep Ensembles. in Advances in Neural Information Processing Systems vol. 30 (Curran Associates, Inc., 2017).

29. Reed, K. S. M. et al. Temporal analysis suggests a reciprocal relationship between 3D chromatin structure and transcription. Cell Rep 41, 111567 (2022).

30. Shrikumar, A., Greenside, P. & Kundaje, A. Learning important features through propagating activation differences. in Proceedings of the 34th International Conference on Machine Learning - Volume 70 3145–3153 (JMLR.org, Sydney, NSW, Australia, 2017).

31. Lieberman-Aiden, E. et al. Comprehensive mapping of long-range interactions reveals folding principles of the human genome. Science 326, 289–293 (2009).

32. Valton, A.-L. & Dekker, J. TAD disruption as oncogenic driver. Curr Opin Genet Dev 36, 34–40 (2016).

33. Hnisz, D. et al. Activation of proto-oncogenes by disruption of chromosome neighborhoods. Science 351, 1454–1458 (2016).

34. Kloetgen, A. et al. Three-dimensional chromatin landscapes in T cell acute lymphoblastic leukemia. Nat Genet 52, 388–400 (2020).

35. Liu, N. Q. et al. The non-coding variant rs1800734 enhances DCLK3 expression through long-range interaction and promotes colorectal cancer progression. Nat Commun 8, 14418 (2017).

36. ENCODE Project Consortium et al. Expanded encyclopaedias of DNA elements in the human and mouse genomes. Nature 583, 699–710 (2020).

37. Tang, F., Yang, Z., Tan, Y. & Li, Y. Super-enhancer function and its application in cancer targeted therapy. NPJ Precis Oncol 4, 2 (2020).

38. Khan, A. & Zhang, X. dbSUPER: a database of super-enhancers in mouse and human genome. Nucleic Acids Res 44, D164–171 (2016).

39. Sahin, M. et al. HiC-DC+ enables systematic 3D interaction calls and differential analysis for Hi-C and HiChIP. Nat Commun 12, 3366 (2021).

40. Yates, J. et al. Cell states and neighborhoods in distinct clinical stages of primary and metastatic esophageal adenocarcinoma. bioRxiv 2024.08.17.608386 (2024) doi:10.1101/2024.08.17.608386.

41. Celià-Terrassa, T. et al. Normal and cancerous mammary stem cells evade interferon-induced constraint through the miR-199a-LCOR axis. Nat Cell Biol 19, 711–723 (2017).

42. Pérez-Núñez, I. et al. LCOR mediates interferon-independent tumor immunogenicity and responsiveness to immune-checkpoint blockade in triple-negative breast cancer. Nat Cancer 3, 355– 370 (2022).

43. Bleu, M. et al. PAX8 and MECOM are interaction partners driving ovarian cancer. Nat Commun 12, 2442 (2021).

44. Ma, Y. et al. CRISPR-mediated MECOM depletion retards tumor growth by reducing cancer stem cell properties in lung squamous cell carcinoma. Mol Ther 30, 3341–3357 (2022).

45. Lou, M. et al. MECOM and the PRDM gene family in uterine endometrial cancer: bioinformatics and experimental insights into pathogenesis and therapeutic potentials. Mol Med 30, 190 (2024).

46. Madry, A., Makelov, A., Schmidt, L., Tsipras, D. & Vladu, A. Towards Deep Learning Models Resistant to Adversarial Attacks. Preprint at 10.48550/arXiv.1706.06083 (2019).

47. Shafahi, A. et al. Adversarial training for free! in Advances in Neural Information Processing Systems vol. 32 (Curran Associates, Inc., 2019).

48. Bai, T., Luo, J., Zhao, J., Wen, B. & Wang, Q. Recent Advances in Adversarial Training for Adversarial Robustness. in vol. 5 4312–4321 (2021).

49. Reiff, S. B. et al. The 4D Nucleome Data Portal as a resource for searching and visualizing curated nucleomics data. Nat Commun 13, 2365 (2022).

50. Imakaev, M. et al. Iterative correction of Hi-C data reveals hallmarks of chromosome organization. Nat Methods 9, 999–1003 (2012).

51. Wolff, J. et al. Galaxy HiCExplorer 3: a web server for reproducible Hi-C, capture Hi-C and single-cell Hi-C data analysis, quality control and visualization. Nucleic Acids Res 48, W177–W184 (2020).

52. Virtanen, P. et al. SciPy 1.0: fundamental algorithms for scientific computing in Python. Nat Methods 17, 261–272 (2020).

53. Ramírez, F. et al. deepTools2: a next generation web server for deep-sequencing data analysis. Nucleic Acids Res 44, W160–165 (2016).

54. Robinson, M. D. & Oshlack, A. A scaling normalization method for differential expression analysis of RNA-seq data. Genome Biol 11, R25 (2010).

55. Rauluseviciute, I. et al. JASPAR 2024: 20th anniversary of the open-access database of transcription factor binding profiles. Nucleic Acids Res 52, D174–D182 (2024).

56. Swindell, W. R. et al. Meta-profiles of gene expression during aging: limited similarities between mouse and human and an unexpectedly decreased inflammatory signature. PLoS One 7, e33204 (2012).

57. Khalid, M. et al. Empirical Evaluation of Activation Functions in Deep Convolution Neural Network for Facial Expression Recognition. in 2020 43rd International Conference on Telecommunications and Signal Processing (TSP) 204–207 (2020). doi:10.1109/TSP49548.2020.9163446.

58. Yamada, Y., Lindenbaum, O., Negahban, S. & Kluger, Y. Feature Selection using Stochastic Gates. in Proceedings of the 37th International Conference on Machine Learning 10648–10659 (PMLR, 2020).

59. Vaswani, A. et al. Attention is All you Need. in Advances in Neural Information Processing Systems vol. 30 (Curran Associates, Inc., 2017).

60. Carty, M. et al. An integrated model for detecting significant chromatin interactions from high-resolution Hi-C data. Nat Commun 8, 15454 (2017).

61. Nassar, L. R. et al. The UCSC Genome Browser database: 2023 update. Nucleic Acids Res 51, D1188–D1195 (2023).

62. Karimzadeh, M., Ernst, C., Kundaje, A. & Hoffman, M. M. Umap and Bismap: quantifying genome and methylome mappability. Nucleic Acids Res 46, e120 (2018).

63. Ansel, J. et al. PyTorch 2: Faster Machine Learning Through Dynamic Python Bytecode Transformation and Graph Compilation. in Proceedings of the 29th ACM International Conference on Architectural Support for Programming Languages and Operating Systems, Volume 2 929–947 (ACM, La Jolla CA USA, 2024). doi:10.1145/3620665.3640366.

64. Kingma, D. P. & Ba, J. Adam: A Method for Stochastic Optimization. Preprint at 10.48550/arXiv.1412.6980 (2017).

65. Rao, S. S. P. et al. A 3D Map of the Human Genome at Kilobase Resolution Reveals Principles of Chromatin Looping. Cell 159, 1665–1680 (2014).

66. Bailey, S. D. et al. ZNF143 provides sequence specificity to secure chromatin interactions at gene promoters. Nat Commun 2, 6186 (2015).

67. Zhou, Q. et al. ZNF143 mediates CTCF-bound promoter-enhancer loops required for murine hematopoietic stem and progenitor cell function. Nat Commun 12, 43 (2021).

68. Magnitov, M. D. et al. ZNF143 is a transcriptional regulator of nuclear-encoded mitochondrial genes that acts independently of looping and CTCF. Mol Cell S1097-2765(24)00956–0 (2024) doi:10.1016/j.molcel.2024.11.031.

69. Loshchilov, I. & Hutter, F. Decoupled Weight Decay Regularization. Preprint at 10.48550/arXiv.1711.05101 (2019).

70. Kelley, D. R. et al. Sequential regulatory activity prediction across chromosomes with convolutional neural networks. Genome Res 28, 739–750 (2018).

71. Dekker, J. et al. The 4D nucleome project. Nature 549, 219–226 (2017).

72. Li, H. et al. The Sequence Alignment/Map format and SAMtools. Bioinformatics 25, 2078–2079 (2009).

73. Lee, J. et al. kundajelab/atac_dnase_pipelines: 0.3.3. Zenodo 10.5281/ZENODO.596029 (2016).

74. Loubiere, V., de Almeida, B. P., Pagani, M. & Stark, A. Developmental and housekeeping transcriptional programs display distinct modes of enhancer-enhancer cooperativity in Drosophila. Nat Commun 15, 8584 (2024).

