## Supplementary Figures for "UniversalEPI: robust prediction of cell type-specific and differential chromatin interactions from DNA sequence and chromatin accessibility"

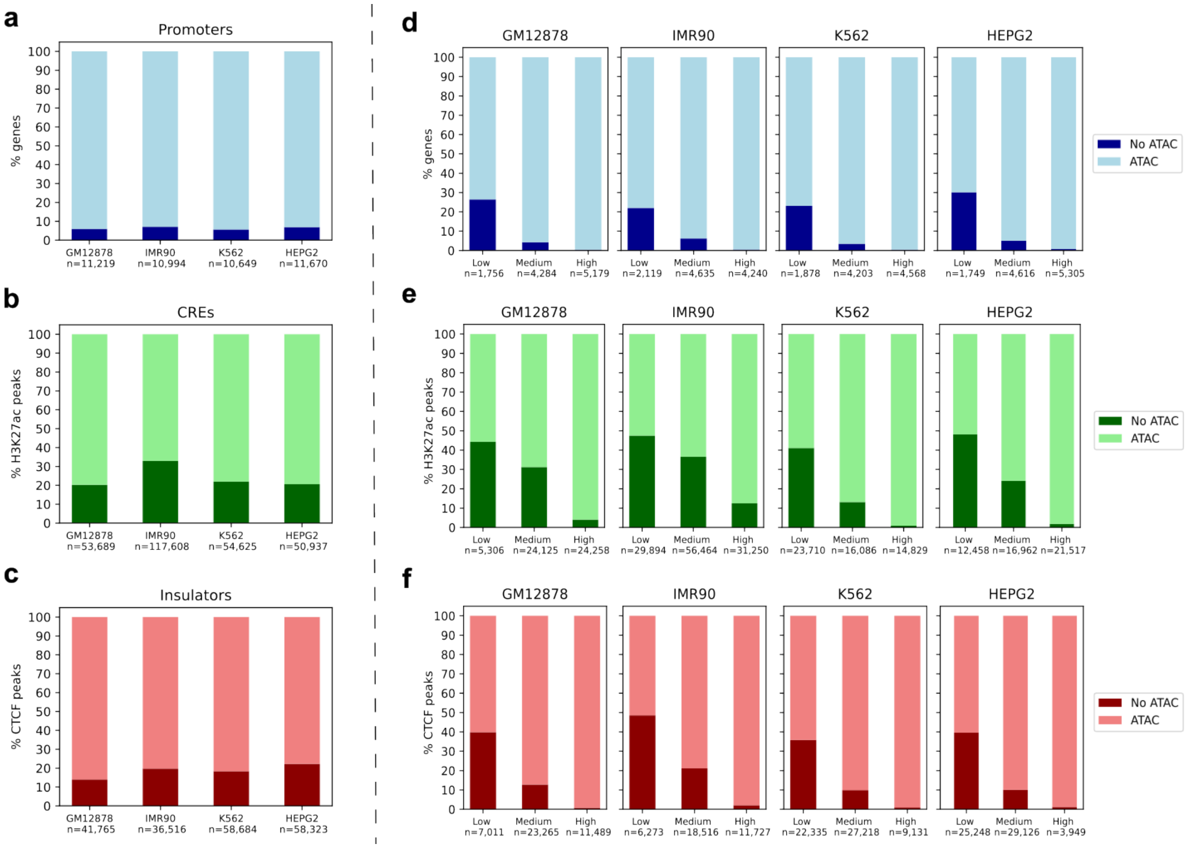


Figure 1: **Information loss by retaining only accessible chromatin regions.** Information lost due to the selection of accessible chromatin regions as measured by the overlap of ATAC-seq peaks with the transcription start site of the **a,** expressed genes (promoter-like), **b,** H3K27ac histone modification (cis-regulatory elements or CREs), and **c,** CTCF sites (insulator-like) in four cell lines: GM12878, IMR90, K562, and HepG2. These cell lines have 186,423, 174,095, 179,240, and 175,487 peaks respectively. Nearly 95% of active promoters overlap with ATAC-seq peaks whereas approximately 80% of active CREs and insulators are captured by ATAC-seq peaks. **d-f,** The promoters, CREs, and insulators not captured by ATAC-seq peaks often have low coverage.

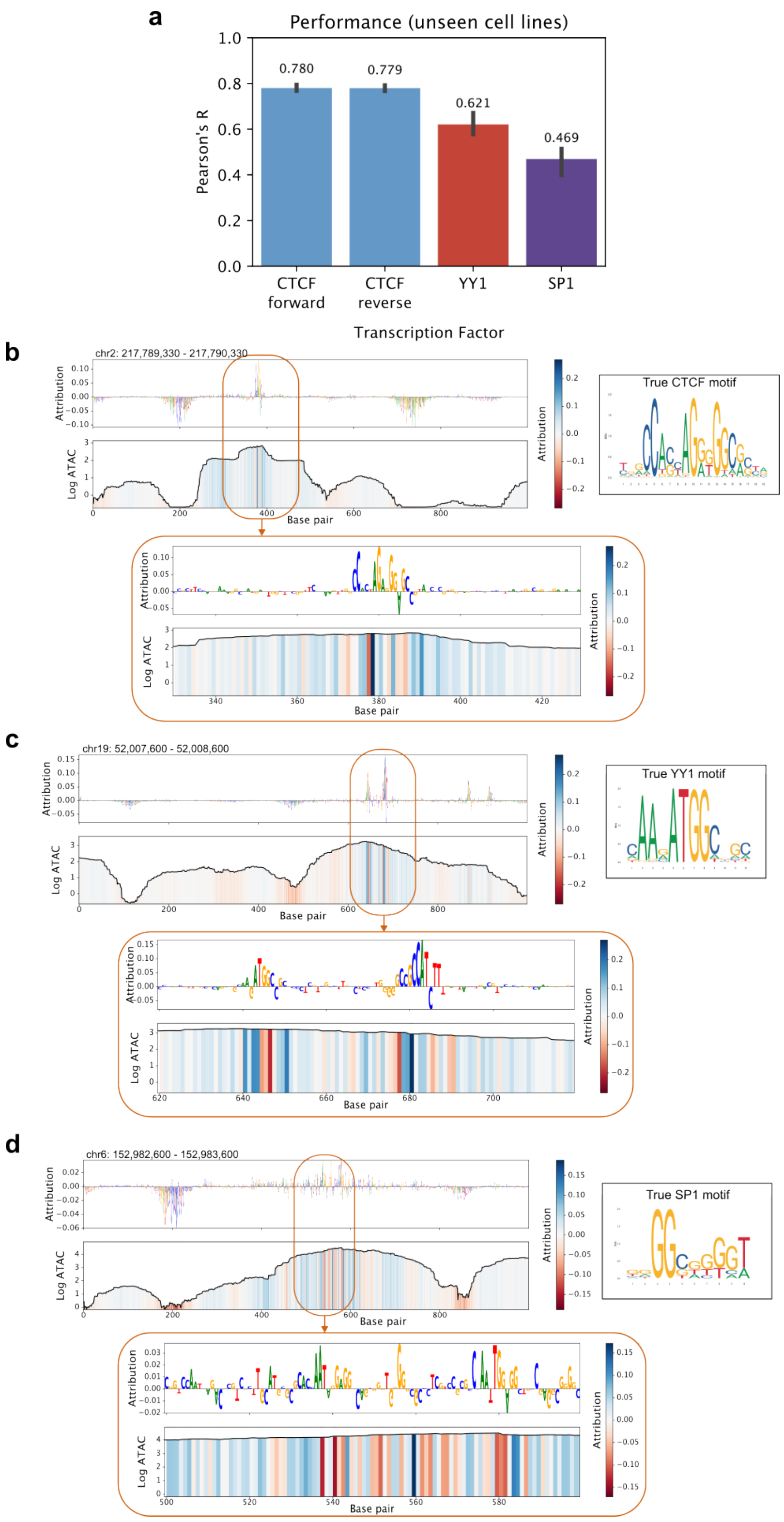


Figure 2: **Performance of first stage of UniversalEPI. a,** Predictive performance of the first stage of UniversalEPI on unseen chromosomes of cell lines that are not seen during training (GM12878, K562, HepG2, A549). The mean score across the four cell lines is reported and the range is represented above the bar. The true motif is obtained from JASPAR. **b,** DeepLIFT attribution scores on unseen chromosome of unseen cell line (HepG2) matches the CTCF motif. **c,** DeepLIFT attribution score is matched with YY1 motif. Two YY1 motifs are identified in this example (one in reverse complement). **d,** SP1 motif is also identified by the model.

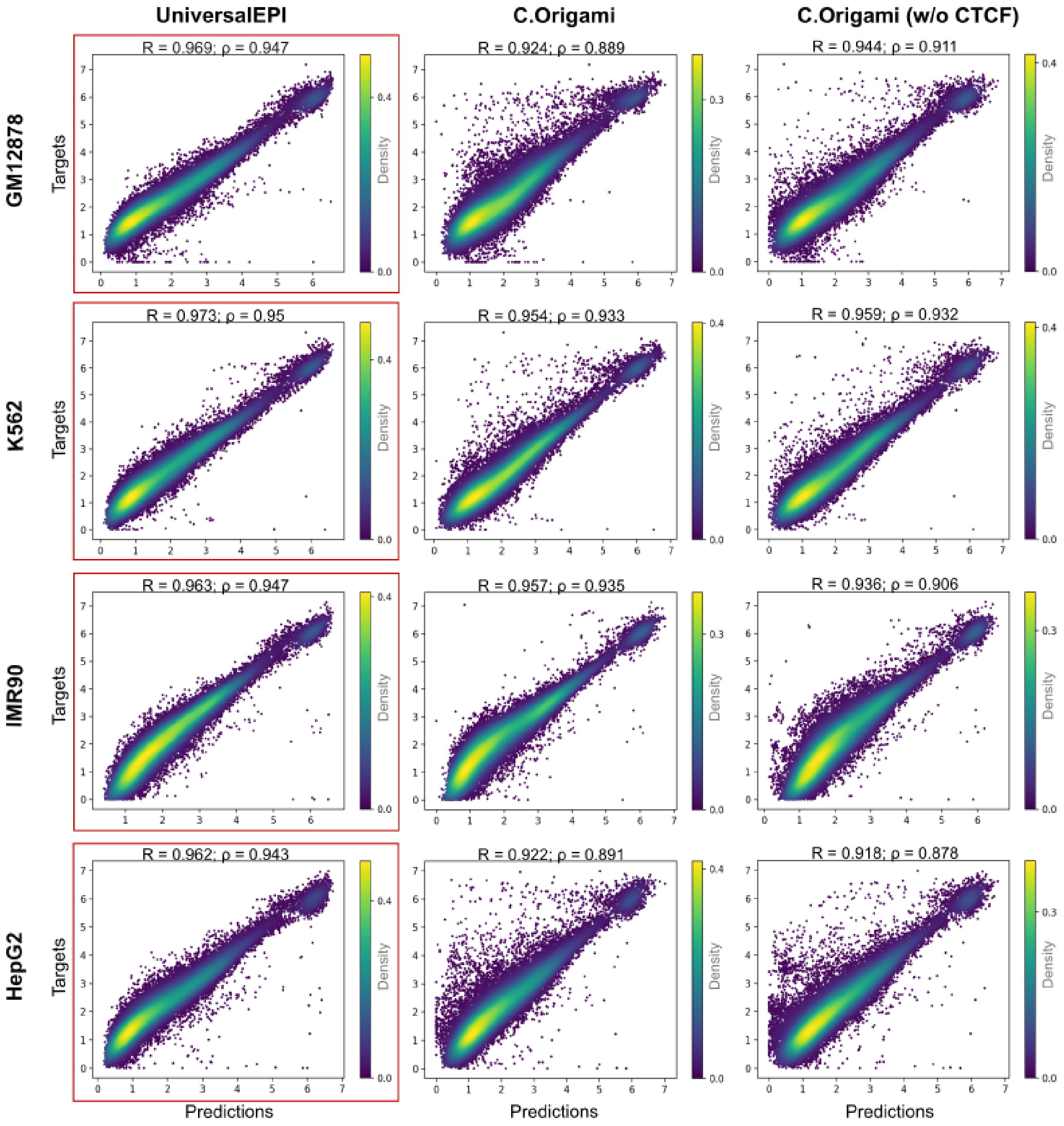


Figure 3: **Performance comparison between UniversalEPI and C.Origami on unseen cell lines.** Comparison of UniversalEPI against state-of-the-art methods like C.Origami [1], and C.Origami (when CTCF ChIP-seq is removed from the input) on the test chromosomes (chromosomes 2, 6, and 19) of cell lines that were not seen by the model during training. Two versions of models are used here: one is trained on GM12878 and K562 cell lines and the other is trained on IMR90 and HepG2 cell lines. Pearson's correlation ($R$) and Spearman's correlation ($\rho$) are calculated for each method and cell line. The best performing method for each cell line (based on $\rho$) is highlighted in red.

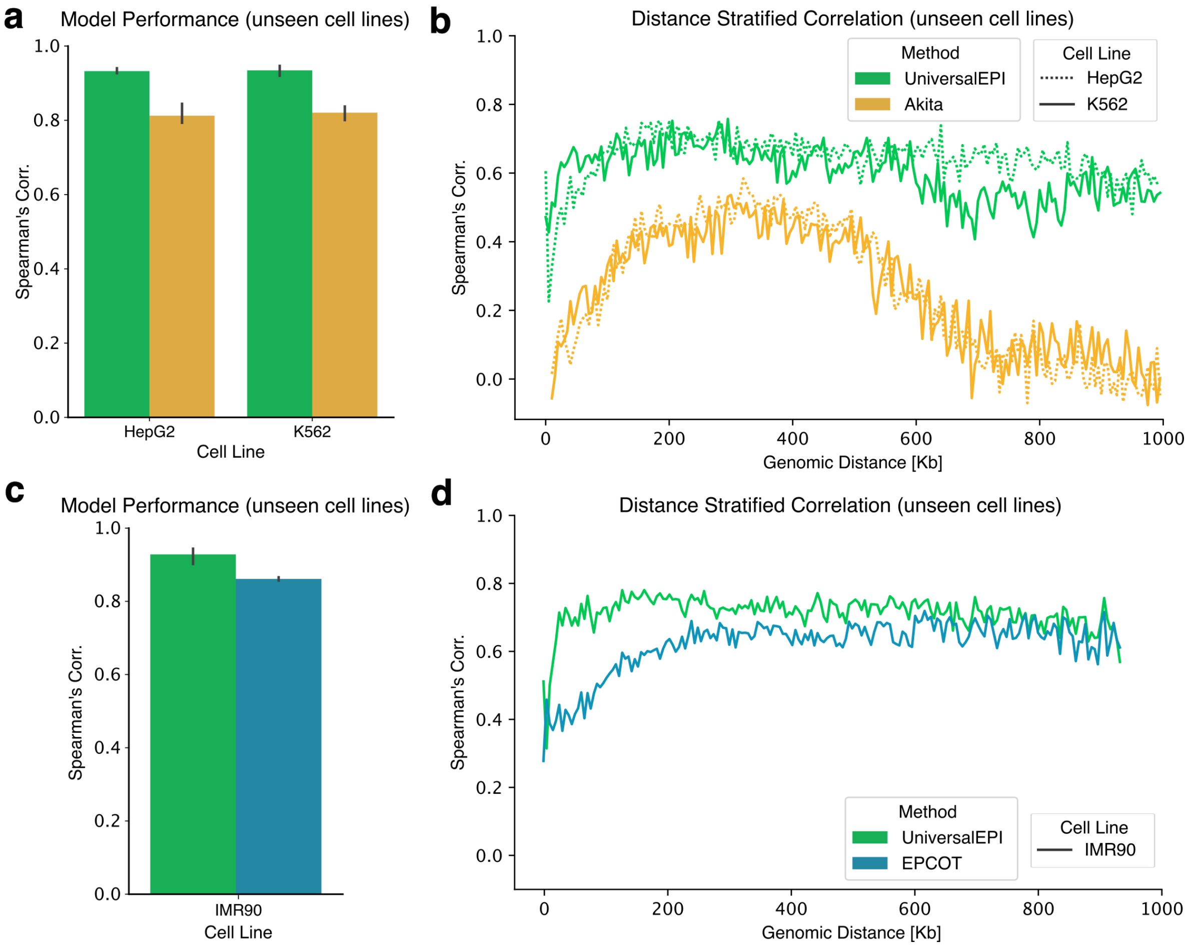


Figure 4: **Performance comparison of UniversalEPI against Akita and EPCOT on unseen cell lines.** **a–b** Comparison between UniversalEPI and Akita [2] on test chromosomes (2, 6, and 19) from previously unseen cell lines (HepG2 and K562). Two UniversalEPI variants are evaluated: one trained on GM12878 and K562, and another on IMR90 and HepG2. **a,** Overall prediction accuracy measured by Spearman correlation. **b,** Performance stratified by genomic distance. **c–d** Comparison between UniversalEPI and EPCOT [3] on the unseen IMR90 cell line. **c,** Overall Spearman correlation. **d,** Distance-stratified performance.

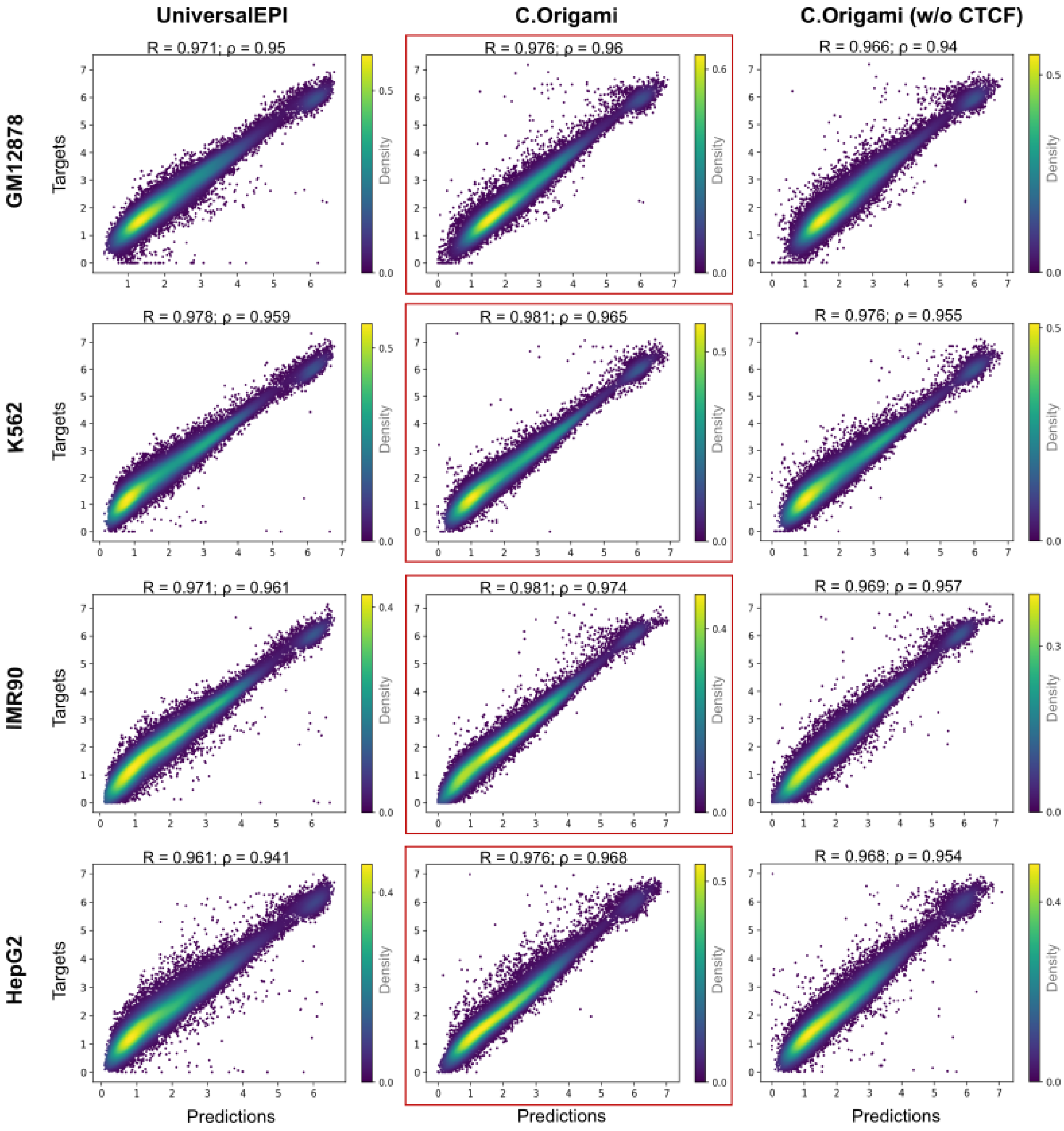


Figure 5: **Performance comparison between UniversalEPI and C.Origami on seen cell lines.** Comparison of UniversalEPI against state-of-the-art methods like C.Origami, and C.Origami (when CTCF ChIP-seq is removed from the input) on the test chromosomes (chr2, chr6, and chr19) of cell lines that were seen by the model during training. Two versions of models are used here: one is trained on GM12878 and K562 cell lines and the other is trained on IMR90 and HepG2 cell lines. Pearson's correlation ($R$) and Spearman's correlation ($\rho$) are calculated for each method and cell line. The best performing method for each cell line (based on $\rho$) is highlighted in red.

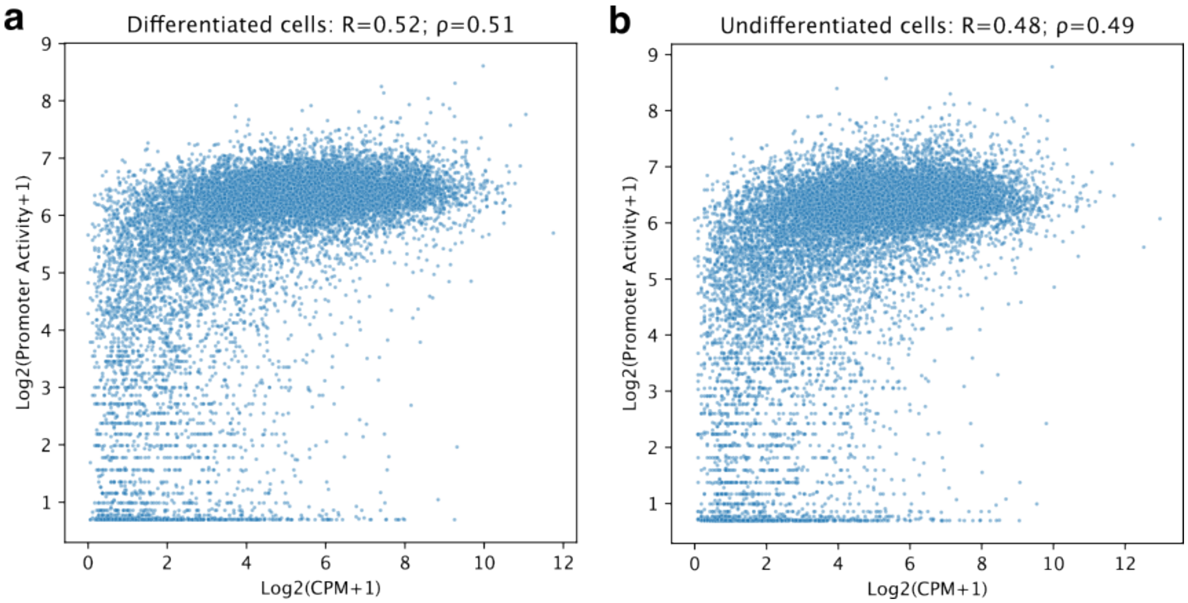


Figure 6: **Correlation of promoter activity with gene expression in EAC. a-b,** Pearson's and Spearman's correlations ($R$ and $\rho$ respectively) of the estimated promoter activity using experimental ATAC-seq and Hi-C derived from UniversalEPI in differentiated and undifferentiated cells in EAC patients.

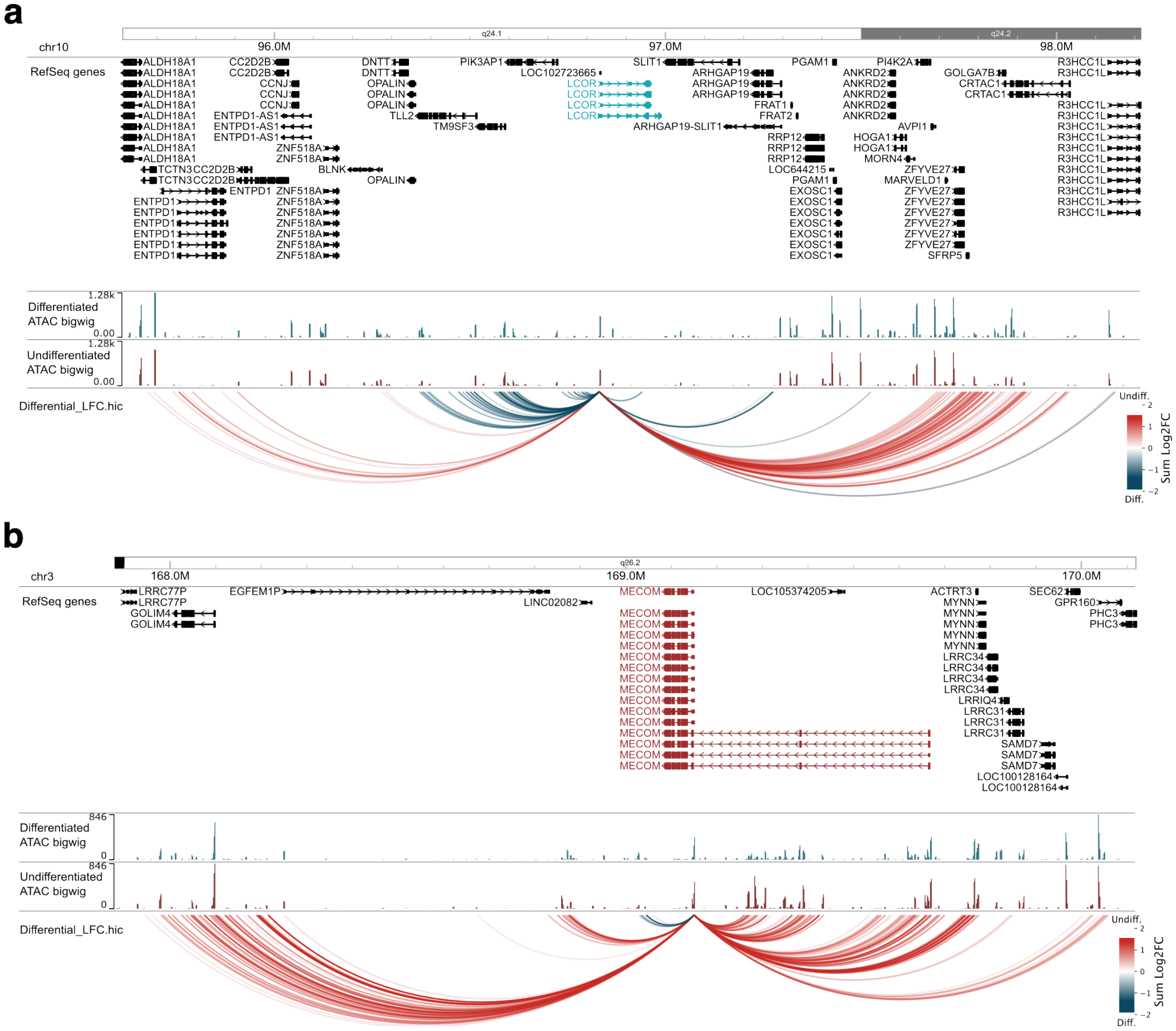


Figure 7: **Differential interactions in EAC without using uncertainty.** Differential interactions between undifferentiated and differentiated cells in EAC are predicted using UniversalEPI. No filtering is applied based on estimated uncertainty. **a,** Different sets of strong interactions are identified in favor of differentiated and undifferentiated cells with LCOR promoter. LCOR is a master transcription factor in differentiated cells. **b,** Similar interactions are shown for MECOM, which is a master transcription factor in undifferentiated cells. The figure is generated using the WashU Epigenome Browser [4].

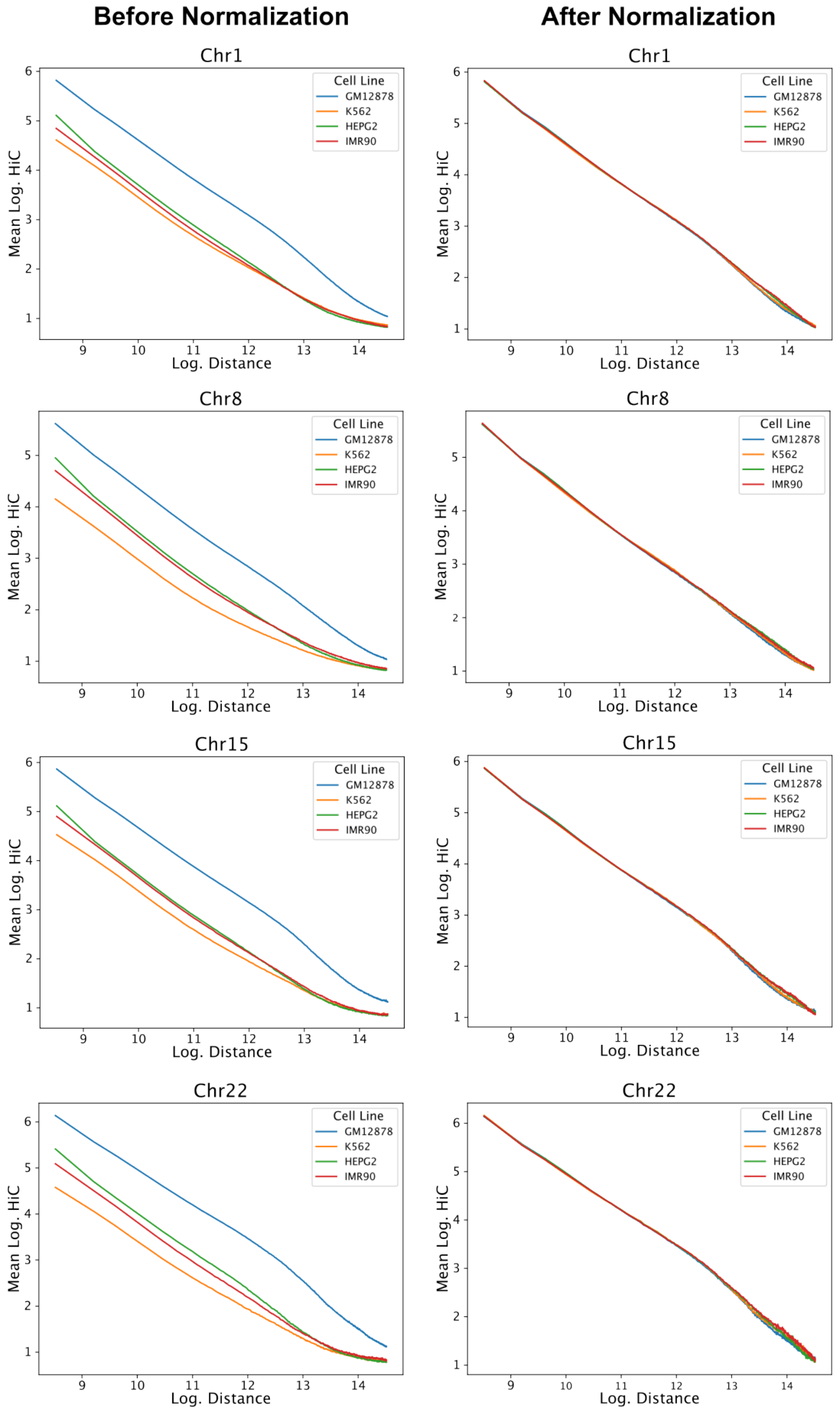


Figure 8: **Cross-cell-type normalization of Hi-C.** Before and after distance-stratified robust z-score normalization on four randomly selected chromosomes (chromosomes 1, 8, 15, and 22) on all cell lines. GM12878 is used as the reference cell line due to its high sequencing depth.

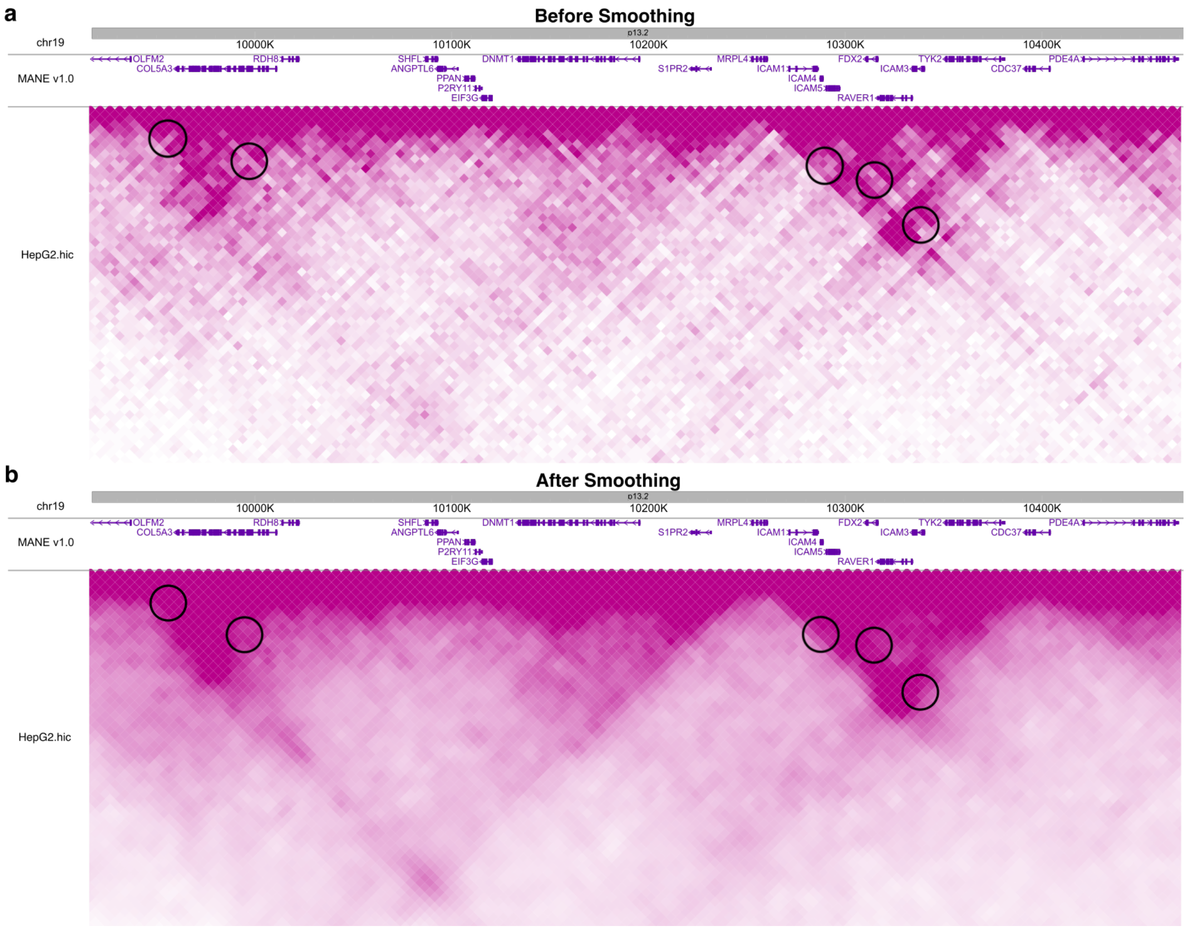


Figure 9: **Smoothing of Hi-C matrices.** The effect of our variable Gaussian smoothing on a randomly selected region of chromosome 19 in the HepG2 cell line. **a,** We can observe gaps (circled) in the original Hi-C matrix that can be attributed to the technical effects. **b,** These gaps are filled using our variable smoothing thereby giving a more realistic Hi-C matrix. The figure is generated using the WashU Epigenome Browser [4].

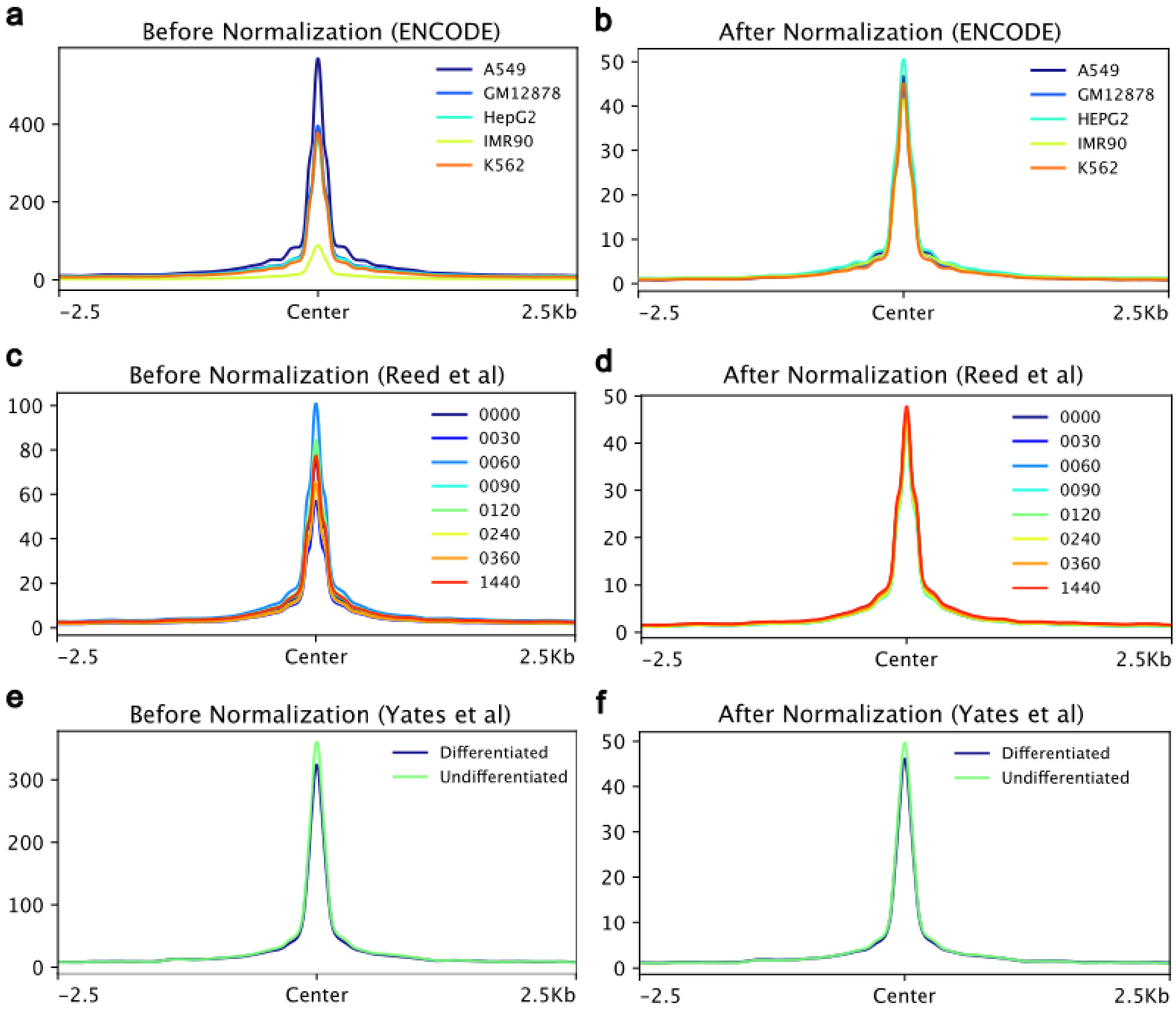


Figure 10: **Cross-cell-type normalization of ATAC-seq signal. a-b,** Before and after applying edgeR's trimmed mean of M-values normalization [5] on ENCODE cell lines [6] with GM12878 as the reference cell line. The conserved active CTCF sites are used as reference regions. **c-d,** Same normalization is applied to the macrophage differentiation data (Reed *et al* [7]) with GM12878 as reference. **e-f,** The ATAC-seq from esophageal carcinoma (Yates *et al* [8]) is normalized using the same normalization with GM12878 as reference.

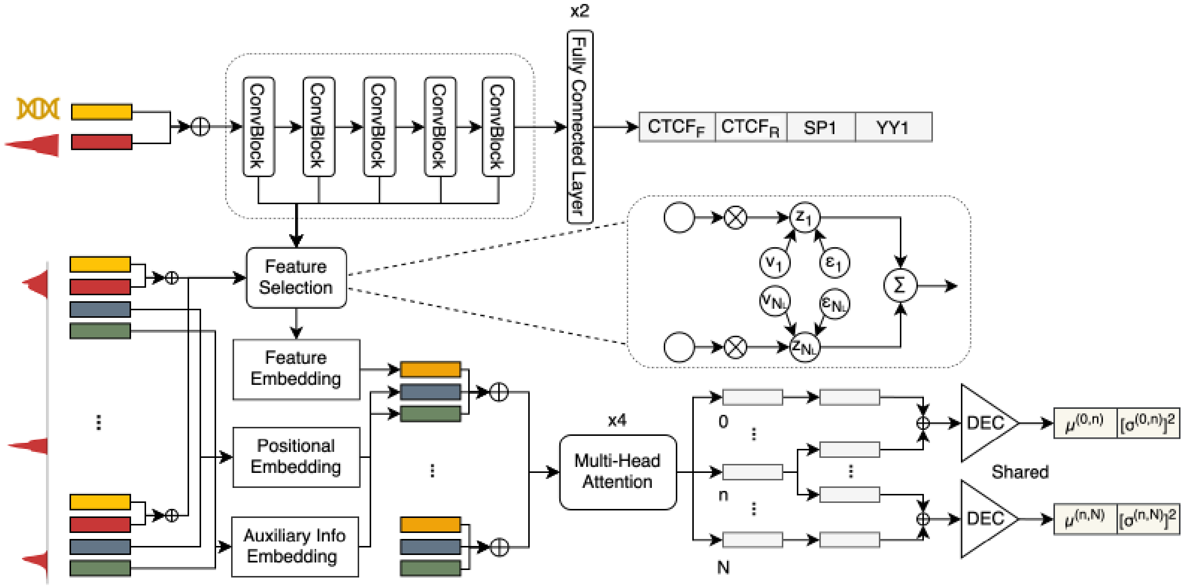


Figure 11: **A graphical illustration of UniversalEPI**. UniversalEPI consists of a CNN based representation network, a feature selection component leveraging stochastic gating (STG), and a transformer based Hi-C prediction network. The representation network takes one-hot encoded DNA sequences and ATAC signals as input, and predicts the binding affinity of four transcription factors. It comprises five convolutional blocks, followed by two fully connected layers. The Hi-C prediction network takes a sequence of ATAC peak regions. Each region is representation by the features from learned convolutions selected using the STG mechanism, its genomic distance and auxilliary information, such as mappability. Before being fed into the multi-head attention layers, each component is embedded via a linear projection layer. The decoder (DEC) operates on pair of output tokens at positions $i$ and $j$ of the attention layers, to estimate the corresponding mean $\mu^{(i,j)}$ and variance $[\sigma^{(i,j)}]^2$ of the Hi-C interaction value. $\bigoplus$ and $\bigotimes$ denote concatenation and multiplication, respectively.

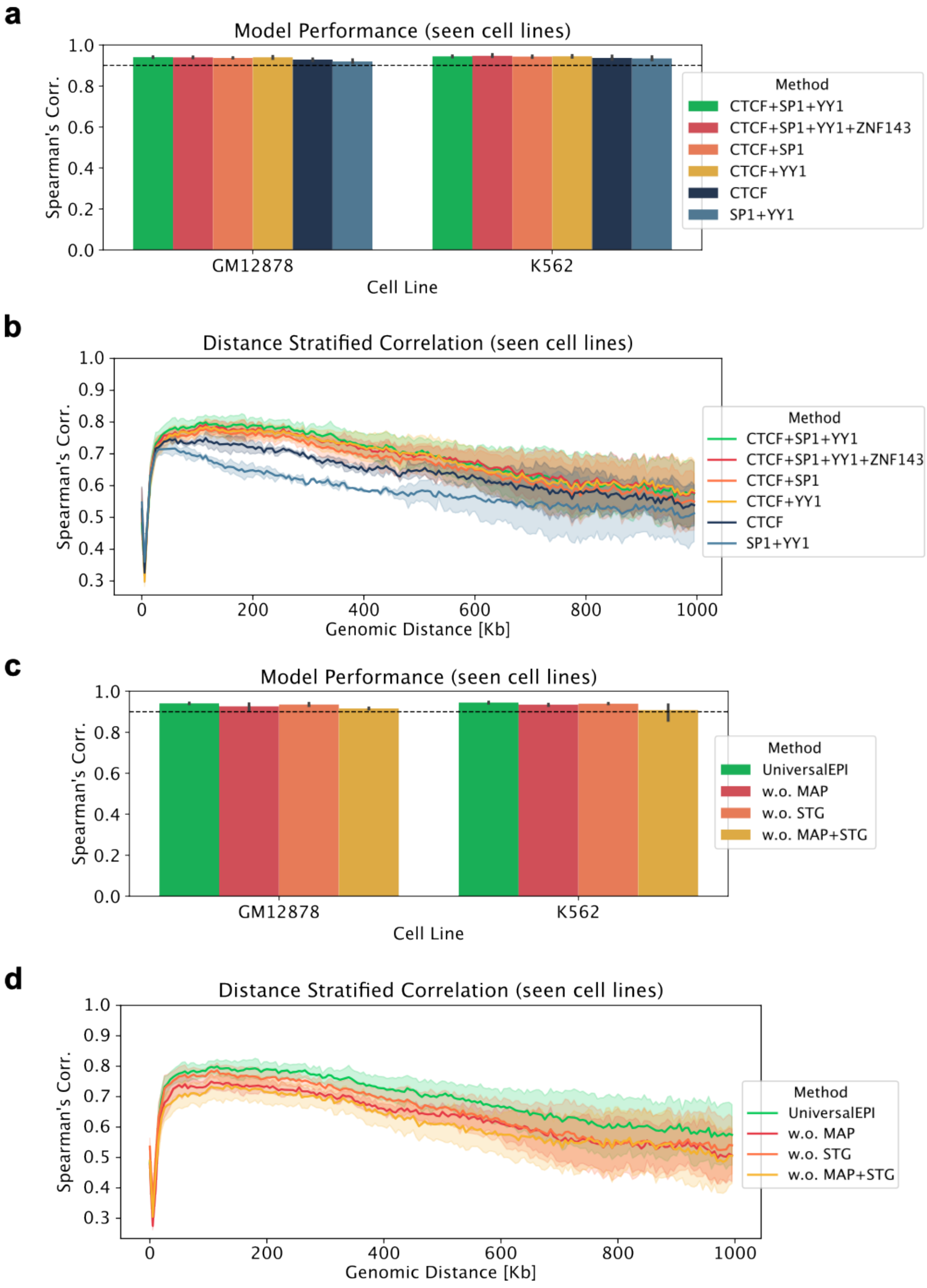


Figure 12: **Ablation studies are performed on UniversalEPI.** Unseen chromosomes of the training cell lines (GM12878 and K562) are used to perform all the ablations. Moreover, the ablations are performed on UniversalEPI without uncertainty estimation and ATAC-seq as input. **a,** Different sets of transcription factors that are used for training the first stage of UniversalEPI are compared based on Spearman's correlation. **b,** Distance-stratified Spearman's correlation is used to reflect differences between models that have similar overall performance. The shaded region represents the variation across the two cell lines. **c,** The effect of using mappability tracks as auxiliary information in the second stage of UniversalEPI and the effect of applying stochastic gating is compared. **d,** Distance-stratified Spearman's correlation is studied as above.

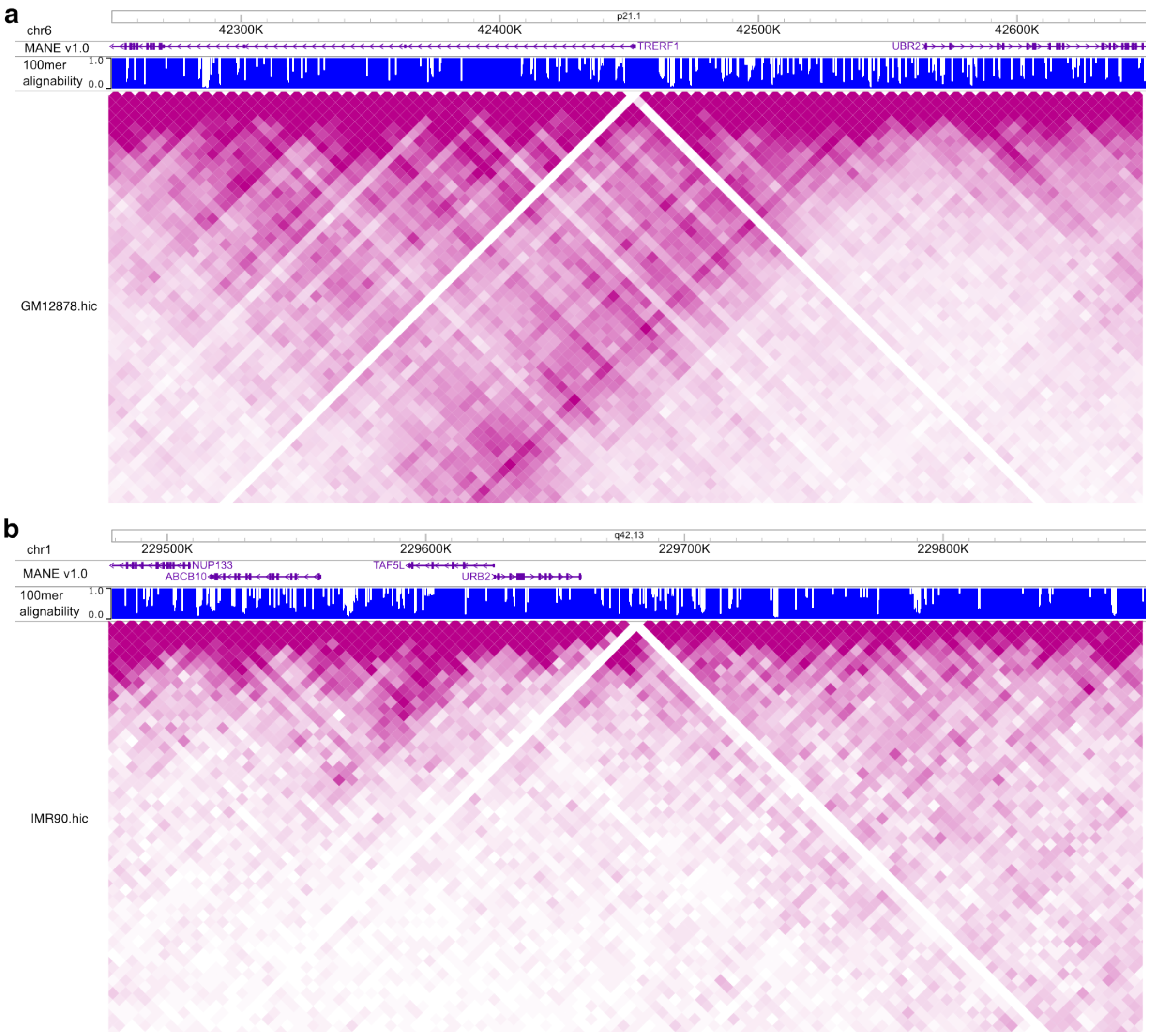


Figure 13: **Examples of blacklisted regions.** Several bins in the Hi-C data of each cell line arbitrarily has no interactions despite being mappable. **a,** An example of such a region in the GM12878 cell line is observed at chromosome 6 for the bin 42,450,000-42,455,000. **b,** Another example can be seen in the IMR90 cell line at chromosome 1 between 229,680,000 and 229,685,000. The figure is generated using the WashU Epigenome Browser [4].

### References


[1] Jimin Tan, Nina Shenker-Tauris, Javier Rodriguez-Hernaez, Eric Wang, Theodore Sakellaropoulos, Francesco Boccalatte, Palaniraja Thandapani, Jane Skok, Iannis Aifantis, David Fenyö, et al. Cell-type-specific prediction of 3d chromatin organization enables high-throughput in silico genetic screening. *Nature Biotechnology*, 41(8):1140–1150, 2023.

[2] Geoff Fudenberg, David R Kelley, and Katherine S Pollard. Predicting 3d genome folding from dna sequence with akita. *Nature Methods*, 17(11):1111–1117, 2020.

[3] Zhenhao Zhang, Fan Feng, Yiyang Qiu, and Jie Liu. A generalizable framework to comprehensively predict epigenome, chromatin organization, and transcriptome. *Nucleic Acids Research*, 51(12):5931–5947, 2023.

[4] Daofeng Li, Silas Hsu, Deepak Purushotham, Renee L Sears, and Ting Wang. Washu epigenome browser update 2019. *Nucleic Acids Research*, 47(W1):W158–W165, 2019.

[5] Mark D Robinson and Alicia Oshlack. A scaling normalization method for differential expression analysis of rna-seq data. *Genome Biology*, 11:1–9, 2010.

[6] Yunhai Luo, Benjamin C Hitz, Idan Gabdank, Jason A Hilton, Meenakshi S Kagda, Bonita Lam, Zachary Myers, Paul Sud, Jennifer Jou, Khine Lin, et al. New developments on the encyclopedia of dna elements (encode) data portal. *Nucleic Acids Research*, 48(D1):D882–D889, 2020.

[7] Kathleen SM Reed, Eric S Davis, Marielle L Bond, Alan Cabrera, Eliza Thulson, Ivana Yoseli Quiroga, Shannon Cassel, Kamisha T Woolery, Isaac Hilton, Hyejung Won, et al. Temporal analysis suggests a reciprocal relationship between 3d chromatin structure and transcription. *Cell Reports*, 41(5), 2022.

[8] Josephine Yates, Camille Mathey-Andrews, Jihye Park, Amanda Garza, Andréanne Gagné, Samantha Hoffman, Kevin Bi, Breanna Titchen, Connor Hennessey, Joshua Remland, et al. Cell states and neighborhoods in distinct clinical stages of primary and metastatic esophageal adenocarcinoma. *Cell Reports Medicine*, 2025.
